# NGF-p75 signaling coordinates skeletal cell migration during bone repair

**DOI:** 10.1101/2021.07.07.451468

**Authors:** Jiajia Xu, Zhao Li, Robert J. Tower, Stefano Negri, Yiyun Wang, Carolyn A. Meyers, Takashi Sono, Qizhi Qin, Amy Lu, Xin Xing, Edward F. McCarthy, Thomas L. Clemens, Aaron W. James

## Abstract

Bone regeneration following injury is initiated by inflammatory signals and occurs in association with infiltration by sensory nerve fibers. Together, these events are believed to coordinate angiogenesis and tissue reprogramming, but the mechanism of coupling immune signals to re-innervation and osteogenesis is unknown. Here, we found that NGF is expressed following cranial bone injury and signals via p75 in resident mesenchymal osteogenic precursors to impact their migration into the damaged tissue. Mice lacking *Ngf* in myeloid cells demonstrated reduced migration of osteogenic precursors to the injury site with consequently delayed bone healing. These features were phenocopied by mice lacking *p75* in *Pdgfra*^+^ osteoblast precursors. Single-cell transcriptomics identified mesenchymal subpopulations with potential roles in cell migration and immune response, altered in the context of *p75* deletion. Together, these results identify the role of p75 signaling pathway in coordinating skeletal cell migration during early bone repair.

## Introduction

The regeneration of craniofacial bones of the mammalian skeleton requires the action of both intrinsic and extrinsic inductive factors from multiple cell types. Unlike the appendicular skeleton, intramembranous cranial bones form, and are healed, without a cartilaginous template by condensations of mesenchymal progenitor cells (*1*). Recent studies from our laboratory have demonstrated an essential role for skeletal sensory nerves in healing of experimental injuries in adult mouse bone, including calvarial bones (*2, 3*). Nerve growth factor (NGF), via its high-affinity receptor tropomyosin receptor kinase A (TrkA), induces skeletal re-innervation, which is essential for later elements of bone repair, including re-vascularization and bone matrix deposition (*2–4*). However, emerging evidence suggests that neurotrophin signaling may have more pleiotropic effects in bone repair than previously considered.

In addition to induction of nerve ingrowth and promotion of nerve survival, pro-NGF binds to its low-affinity receptor p75 (encoded by *Ngfr*), which is present in a variety of mesenchymal cells (*5*). p75 expression has been correlated to cell migration and invasion in the fields of developmental and cancer biology. For example, Schwann cell migration from the dorsal root ganglia is significantly impaired in p75 null embryos (*6*). Likewise, p75-dependent signaling induces the migration of melanoma cells (*7*). Similar observations have been made in neoplastic cells, in which stable knockdown of p75 in melanoma cells reduced cell migration and expression of major regulatory gene networks for cell migration (*8, 9*). NGF binding to the p75 receptor stimulates migration and initiates recruitment of various adaptors, which activate nuclear factor kappa B (NF-κB), Ras homolog family member a (RhoA), and c-Jun N-terminal kinase (JNK) signaling (*10, 11*).

In this study, we have identified an alternate role for NGF signaling in the bone reparative process, independent of skeletal sensory nerves. Depletion of *p75* in platelet-derived growth factor receptor α (PDGFRα)^+^ mesenchymal cells impaired cellular migration into an osseous wound. Transcriptomics identified mesenchymal subpopulations with potential roles in cell migration and immune response, altered in the context of *p75* deletion. Finally, we identified a new mechanism for NGF-expressing macrophages which stimulate skeletal cell migration during early bone repair.

## Results

### Ngf^LysM^ animals display defective bone repair associated with reduced stromal cell migration

In a previous study, we showed that NGF expression is acutely upregulated in both monocytes/macrophages and in resident mesenchymal cells (*3*). To determine the contribution of monocyte/macrophage-derived NGF to cranial bone repair, *Ngf*^LysM^ mice were subjected to frontal bone injuries and compared to *Ngf*^fl/fl^ control mice (**Fig. 1A**). Micro-computed tomography (Micro-CT) reconstructions and cross-sectional images within the defect site demonstrated impaired re-ossification among *Ngf*^LysM^ mice in comparison to *Ngf*^fl/fl^ mice (**Fig. 1B**). In order to investigate a potential link between macrophage-derived NGF and stromal cells within the injured tissue, cellular migration was assessed *in vitro* and *in vivo* using *Ngf*^LysM^ or *Ngf*^fl/fl^ animals (**Fig. 1C-J**). Deletion of *Ngf* in LysM^+^ monocytes/macrophages led to a prominent decrease in stromal cell presence within the mid-defect site, as assessed by PDGFRα immunostaining (**Fig. 1C**), while no change in rates of cellular proliferation or apoptosis was observed, assessed by Ki67 and TUNEL staining, respectively (**Fig. 1D,E**). Flow cytometry for PDGFRα was performed after microdissection bone defect site, which again showed a prominent reduction in PDGFRα^+^ cell frequency within *Ngf*^LysM^ defect sites (**Fig. 1F,G, Table S1**). To directly address the potential paracrine effects of macrophages on calvarial stromal cells, conditioned medium (CM) experiments were performed, harvesting CM from either *Ngf*^LysM^ or *Ngf*^fl/fl^ macrophages (**Fig. 1H,I**). Results demonstrated that CM from macrophages derived from *Ngf*^fl/fl^ mice promoted neonatal mouse calvarial cell (NMCC) migration assessed by scratch wound healing assay, but this pro-migratory effect was not seen in NGF-depleted CM from *Ngf*^LysM^ macrophages (**Fig. 1J**). A similar result was observed with invasion assays. NMCCs were placed on the membrane of the upper chamber, while the CM was added to the bottom well. After 4 h, cells that moved through the membrane were stained. Macrophage CM derived from *Ngf*^fl/fl^, but not *Ngf*^LysM^, induced transwell invasion (**Fig. 1K**). Next, the effects of macrophage CM were assayed on NMCCs derived from conditional gene deletion of *p75* animals (**Fig. 1L,M**). After *in vitro* exposure to 4-Hydroxytamoxifen, a significant reduction in *p75* expression was confirmed among *p75*^PDGFRα^ NMCCs (**Fig. 1L**). Without CM, *p75* knockout in NMCCs was observed to inhibit cell migration. Among control cells without 4-Hydroxytamoxifen treatment, macrophage CM again induced cellular migration. Notably, induction of cellular migration by macrophage CM was not seen among *p75* knockout calvarial cells (**Fig. 1M**). In sum, macrophages stimulate calvarial cell migration in a paracrine fashion, which is dependent on *Ngf* expression. Moreover, expression of NGF’s low-affinity receptor *p75* is similarly required for macrophage-induced cell migration.

**Fig. 1.**
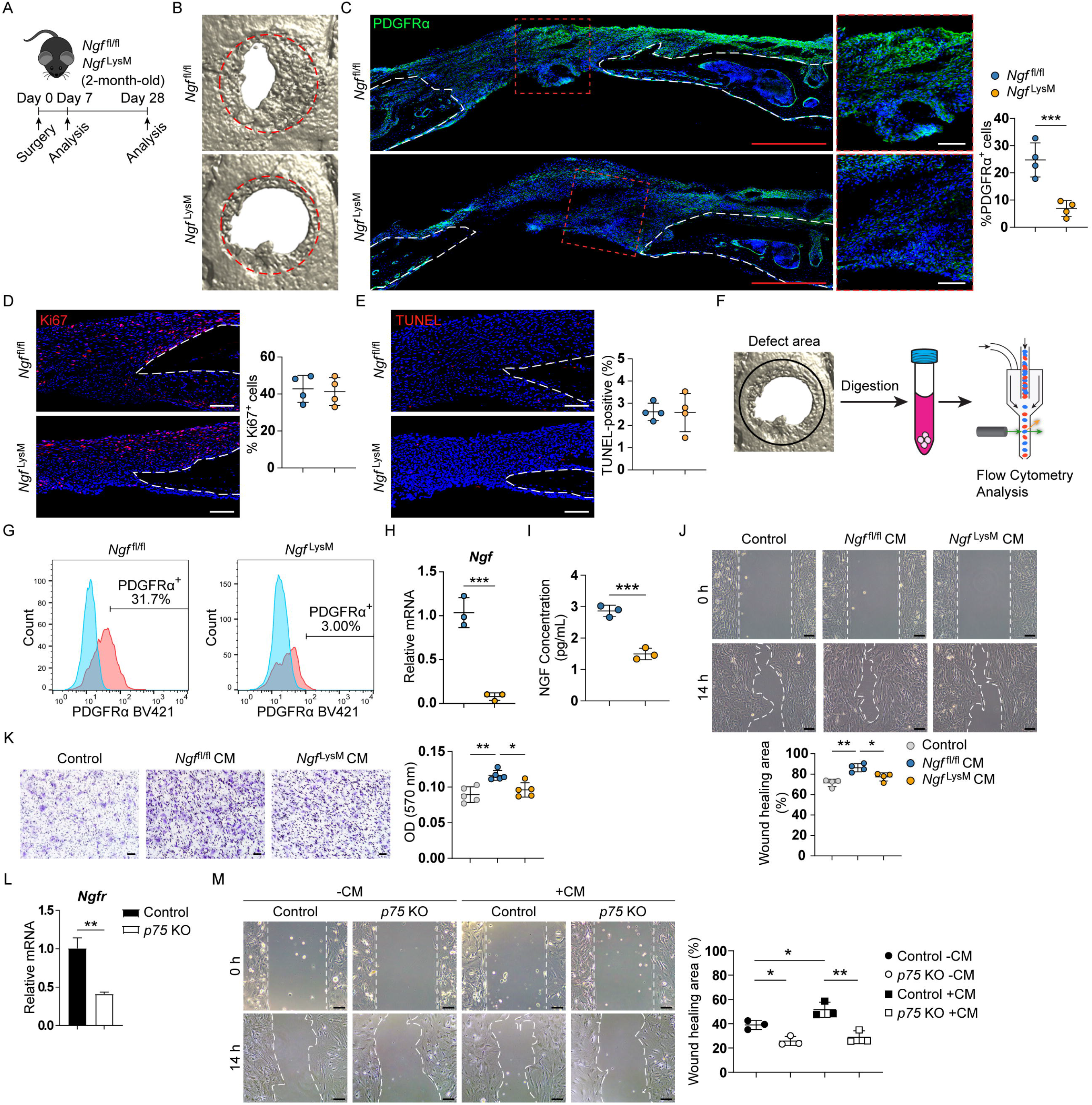
Macrophage-derived NGF positively regulates bone repair and calvarial cell migration. **(A)** Schematic of experiment, in which *Ngf*^fl/fl^ or *Ngf*^LysM^ animals were subjected to frontal bone injury, with analysis performed 7 or 28d post-operatively. **(B)** Micro-CT reconstructions of the defect site at d28 post-injury. Margins of original defect are indicated by dashed red lines. **(C)** PDGFRα^+^ osteoprogenitor cell migration among *Ngf*^fl/fl^ and *Ngf*^LysM^ animals, as assessed by PDGFRα immunofluorescent staining at d7 post-injury. White dashed lines indicate bone edges. **(D,E)** Cellular proliferation **(D)** and apoptosis **(E)** at the bone defect edge at d7 post-injury. **(F)** Schematic of experiment, in which calvarial from *Ngf*^fl/fl^ or *Ngf*^LysM^ animals were harvested at d7 post-injury and only the defect area was left (black circle). After digestion, cells were incubated with antibodies and analyzed by flow cytometry. **(G)** Percentage of PDGFRα^+^ cells among freshly isolated CD31^-^CD45^-^Ter119^-^ cells in defect area. **(H,I)** Validation of *Ngf* deletion in macrophages from *Ngf*^LysM^ mice by RT-PCR (**H**) and ELISA (**I**). **(J,K)** NMCC migration assessed by **(J)** scratch wound healing or **(K)** transwell assay with normal medium (Control) and conditional medium of cultured macrophages from *Ngf*^fl/fl^ or *Ngf*^LysM^ mice. For transwell assay, NMCCs were placed on the membrane of the upper chamber, while medium was added to the bottom well. After 4h, cells were stained. **(L)** Validation of *Ngfr* deletion in NMCCs from *p75*^PDGFRα^ mice after 4-Hydroxytamoxifen treatment. **(M)** Migration of NMCCs treated with or without 4-Hydroxytamoxifen assessed by scratch wound healing assay with normal and conditional medium of cultured macrophages from *Ngf*^fl/fl^ mice. Dot plots represent an individual sample or animal. Data are represented as mean ± 1SD. Scale bar: 500μm (Red) and 100μm (White). N=5 mice **(B)** and 4 mice **(C-G)** per group. \**P*<0.05, \*\**P*<0.01, and \*\*\**P*<0.001 as assessed using a two-tailed Student’s *t*-test or one-way ANOVA.

### p75 expression is required for stromal cell migration and calvarial regeneration

In order to validate our *in vitro* findings, *p75*-expressing cells in the uninjured skull and injured site were identified using a previously validated PDGFRα-CreER^T2^;mT/mG (PDGFRα^mT/mG^) reporter animal (*12*) (**Fig. 2A,B**). Lineage tracing of PDGFRα-expressing cells was based on tamoxifen (TM) administration to promote the activity of inducible Cre, starting at 8 weeks of age. Within the frontal bone, p75 protein was detected at low levels in the overlying periosteum and coincided with PDGFRα reporter activity in bone lining cells (**Fig. 2B**). Following injury, a notable expansion of the p75^+^Pdgfrα^+^ cell population was detected along the outer edge of the injury defect site (**Fig. 2B**). To assess the requirement for p75 in calvarial defect repair, *p75*^fl/fl^ animals were crossed with PDGFRα^mT/mG^ lines to yield *p75*^fl/fl^;PDGFRα^mT/mG^ (*p75*^PDGFRα^) animals. *p75*^fl/fl^;mT/mG animals (*p75*^fl/fl^) were used as controls. Validation for *p75* deletion was performed using p75 immunohistochemistry on calvarial injury sites within the *p75*^fl/fl^ and *p75*^PDGFRα^ mice (**Fig. 2C**). p75 immunohistochemical staining was significantly reduced in *p75*^PDGFRα^ defect sites, in comparison to *p75*^fl/fl^ control defects. Next, frontal bone healing was assessed following *p75* deletion in PDGFRα-expressing cells over a 4-week period (**Fig. 2D-I**). Results demonstrated impaired bone healing among *p75*^PDGFRα^ animals, relative to *p75*^fl/fl^ controls. Micro-CT reconstructions and cross-sectional images demonstrated impaired re-ossification among *p75*^PDGFRα^ mice (**Fig. 2D**). Quantitative micro-CT metrics of bone healing were reduced among *p75*^PDGFRα^ mice, including Bone Volume (BV, **Fig. 2E**, 68.3% reduction), fractional Bone Volume (BV/TV, **Fig. 2F**, 68.3% reduction), mean diameter of the bone defect area (**Fig. 2G**, 40.9% increase), and Bone fractional area (BFA, **Fig. 2H**, 54.8% reduction). H&E staining confirmed a significant impairment of defect healing between bony fronts in *p75* mutant mice (**Fig. 2I**, black arrowheads).

**Fig. 2.**
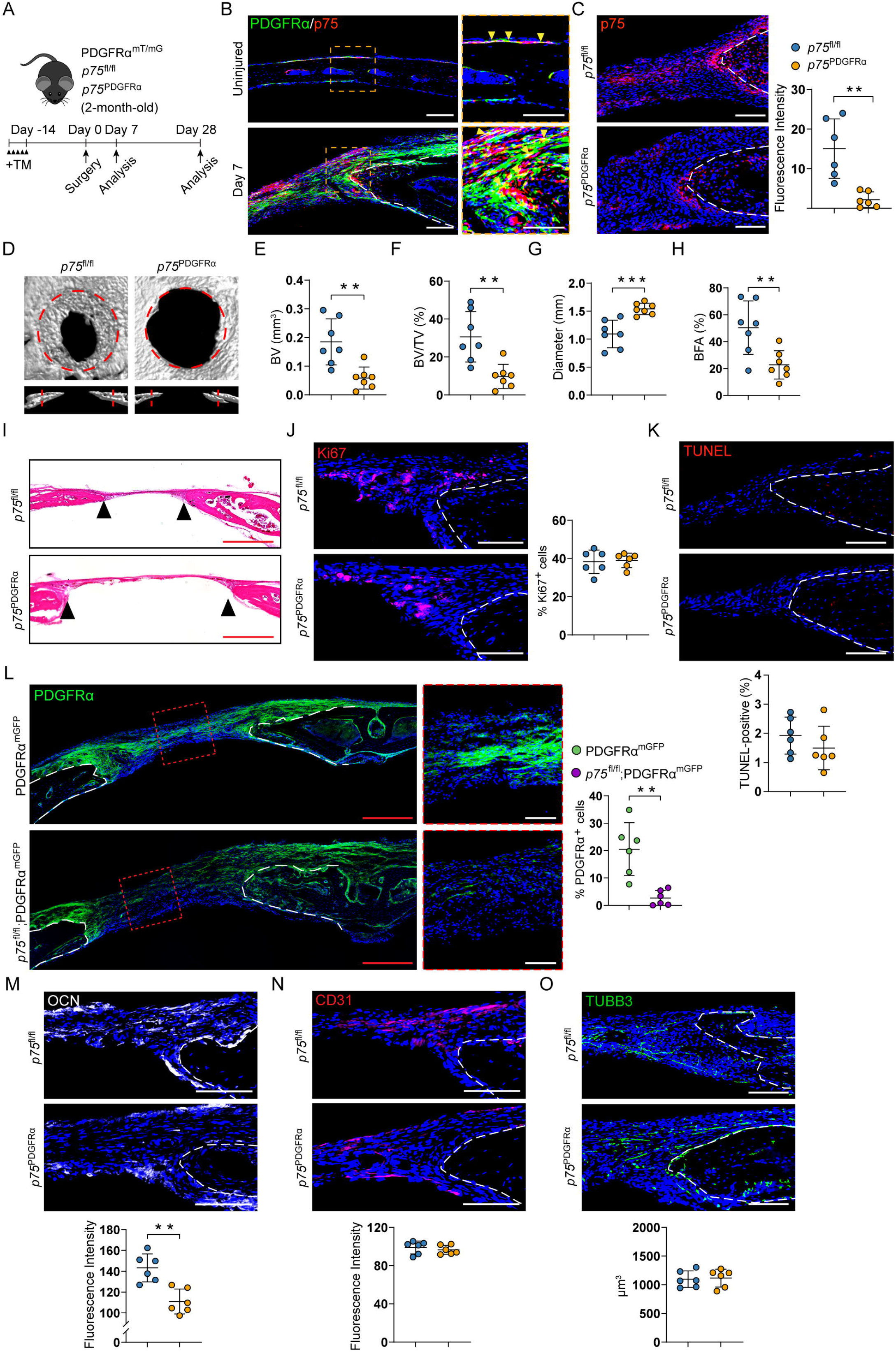
Deletion of *p75* in Pdgfrα-expressing cells impairs stromal cell migration and calvarial bone defect repair. **(A)** Schematic of experiment, in which *p75*^fl/fl^ or *p75*^PDGFRα^ animals were administered tamoxifen (TM), followed by frontal bone injury after 14d, with analysis performed 7 or 28d post-operatively. **(B)** Immunohistochemistry for p75 (red) on the uninjured or calvarial injury site (d7 post-injury) within PDGFRα^mGFP^ reporter sections. Yellow arrowheads indicate p75 and PDGFRα colocalization. TdTomato is not shown. **(C)** Validation of *p75* deletion by p75 immunohistochemistry in either *p75*^fl/fl^ or *p75*^PDGFRα^ mice, d7 post-injury. **(D)** Micro-CT reconstructions of the defect site in a top-down view (above) and coronal cross-sectional images (below) among *p75*^fl/fl^ and *p75*^PDGFRα^ animals, 28d post-injury. Margins of original defect are indicated by dashed red lines. **(E-H)** Micro-CT quantification of bone healing among *p75*^fl/fl^ and *p75*^PDGFRα^ mice (28d post-injury), including **(E)** BV, **(F)** BV/TV, **(G)** residual defect diameter, and **(H)** BFA. **(I)** H&E staining of coronal cross-section of the healing defect site from *p75*^fl/fl^ and *p75*^PDGFRα^ mice at d28 post-injury. Black arrowheads indicate healing bone edges. **(J,K)** Cellular proliferation **(J)** and apoptosis **(K)** at the bone defect edge, as assessed by Ki67 and TUNEL immunofluorescent staining at d7 post-injury. **(L)** Tile scan (left) and high-magnification images of the central defect (right) demonstrate migration of GFP^+^ (PDGFRα^mGFP^) progenitor cells into the defect site at d7 post-injury. Tdtomato is not shown. **(M-O)** Immunohistochemical staining and semi-quantitative analysis of OCN **(M)**, CD31 **(N)**, and TUBB3 **(O)** within the calvarial defect, d28 post-injury. Dashed white lines indicate bone edges. In graphs, each dot represents a single animal. Red scale bar: 500 μm. White scale bar: 100 μm. N=3 mice **(B)** and 6-7 mice **(C-O)** per group. Data are represented as mean ± 1SD. \**P*<0.05, \*\**P*<0.01, and \*\*\**P*<0.001 as assessed using a two-tailed Student’s *t*-test.

To determine potential mechanisms driving this impaired bone healing, Ki67 and TUNEL staining was first conducted to interrogate potential differences in proliferation and apoptosis (**Fig. 2J,K**). These results failed to show any significant differences, suggesting the impaired healing observed in *p75*^PDGFRα^ mice was not the result of reduced cell proliferation or increased apoptosis. Another potential mechanism through which defect repair could be inhibited in p75 mutant mice is the failed recruitment of regenerating mesenchymal cells. To assess progenitor cell migration into the defect, a lineage tracing strategy was employed using PDGFRα^mT/mG^ animals. PDGFRα reporter activity highlights stromal cells which are recruited to populate the bone injury site. Deletion of *p75* in PDGFRα^+^ stromal progenitors resulted in a prominent decrease in stromal cell migration into the central defect region (**Fig. 2L**), also confirmed at an earlier timepoint post-injury (**Fig. S1**). This impaired recruitment of PDGFRα^+^ progenitors led to a reduced number of mature osteoblasts among *p75*^PDGFRα^ animals, as assessed by immunohistochemical staining for Osteocalcin (**Fig. 2M**). Our previous reports have demonstrated that specific impairment of the NGF-TrkA signaling axis leads to an impaired invasion of both the vasculature and sensory nerves into the cranial defect region (*3*). To determine whether a similar phenomenon may also be contributing to our impaired healing deficiencies, vascularity and innervation within the defect site were next evaluated (**Fig. 2N,O**). CD31 immunostaining of endothelium and TUBB3 immunostaining of axons suggested no difference in defect site revascularization or re-innervation among *p75*^fl/fl^ and *p75*^PDGFRα^ mice. In aggregate, inhibition of p75 signaling in PDGFRα^+^ stromal cells led to significant reductions in injury-associated cell migration and a significant delay in bone defect healing, independent of changes in neurovasculature in bone.

### Single-cell transcriptomic profiling of total cells and mesenchymal lineage subpopulation

To understand the mechanisms driving impaired defect repair in *p75*-deficient mice, we employed single-cell RNA sequencing (scRNA-seq) of cells derived from the defect site of *p75*^fl/fl^ and *p75*^PDGFRα^ mice at 7 days post-injury (**Fig. 3A**). Unsupervised clustering identified 8 groups (**Fig. 3B**), identified using expression patterns of known marker genes, including 2 groups of mesenchymal lineage cells (expressing *Paired related homeobox 1* (*Prrx1*)*, Platelet-derived growth factor receptor alpha* (*Pdgfra*)*, Msh homeobox 1* (*Msx1*)*, Twist family BHLH transcription factor 1* (*Twist1*), and *Runt-related transcription factor 2* (*Runx2*)), 5 groups of hematopoietic cells (expressing *Protein tyrosine phosphatase receptor type C* (*Ptprc*) encoding CD45), and 1 group of endothelial cells/pericytes (expressing *Regulator of G-protein signaling 5* (*Rgs5*), and *Endomucin* (*Emcn*)) (**Fig. 3C**). While each cluster was represented by cells from both *p75*^fl/fl^ and *p75*^PDGFRα^ mice, some shifts in cluster proportions were observed, including enrichment of mesenchymal cluster cells derived from *p75*^fl/fl^ mice (**Fig. S2A,B**). Specificity of Cre recombination was confirmed in sequenced data in which EGFP expression was restricted to the PDGFRα-expressing mesenchymal clusters (**Fig. S2C**) and almost exclusively expressed in *p75*^PDGFRα^-derived cells (**Fig. S2D**). Pathway analysis of differentially expressed genes (DEGs) revealed GO term enrichment in cell adhesion, positive regulation of cell migration, osteoblast differentiation, and wound healing in *p75*^fl/fl^ mice (**Fig. 3D**). In contrast, immune system process, inflammatory response, and regulation of cell shape were enriched in cells from *p75*^PDGFRα^ mice, implying altered cell shape, differentiation and matrix interactions (**Fig. 3D**). Focused analysis of mesenchymal lineage cells yielded 8 subpopulations (**Fig. 3E**). Early stem markers *Cd34*, *Thy1* were predominantly expressed in subclusters 1 and 2, while multi-lineage progenitor markers Actin alpha 2 (*Acta2*) and Scleraxis (*Scx*) showed enriched expression in subclusters 3/4 (**Fig. 3F**). Subcluster 7 was classified as being likely derived from the dura mater owing to the selective expression of *Forkhead box protein C2* (*Foxc2*), while subcluster 8 cells were found to express mature osteoblast markers *Bone gamma-carboxyglutamate protein 2* (*Bglap2*) and *Secreted phosphoprotein 1* (*Spp1*) (**Fig. 3F**). Deletion of *p75* in mesenchymal cells was next validated by downstream signaling-related gene expression, including genes of Rho, NF-κB, and mitogen-activated protein kinase (MAPK) signaling pathways (**Fig. S3**). Pathway analysis of the total mesenchymal cell populations showed that osteogenesis-related signaling pathways were inhibited among *p75*^PDGFRα^ cells, including Notch, Bone morphogenetic protein (BMP), Transforming growth factor beta (TGFβ), Fibroblast growth factor (FGF), and Hippo signaling (**Fig. 3G**). Potential defects in cellular migration among *p75*^PDGFRα^ mesenchymal cells were next evaluated. Interestingly, mesenchymal subcluster 5 which expressed high levels of *Itgb1* demonstrated specific enrichment in GO terms such as cell adhesion, regulation of actin cytoskeleton organization, and cell migration (**Fig. 3H**, a complete list of DEGs of subcluster 5 shown in **Table S2**). Owing to our previous results suggesting an impaired migration/recruitment of progenitor cells into the defect area, we analyzed gene lists linked to biological functions needed for cell migration (**Fig. 3I**). Consistent with our cluster DEG assessment and *in vivo* results, cluster 5 showed significant enrichment for genes linked to receptor-mediated extracellular matrix interaction and cytokine receptor-mediated signaling, both of which were reduced in total *p75*^PDGFRα^ cells or subcluster 5 *p75*^PDGFRα^ cells in relation to *p75*^fl/fl^ controls (**Fig. 3I**). This data suggested that p75 signaling within mesenchymal lineage cells is essential for cellular migration and osteogenic differentiation during bone injury repair.

**Fig. 3.**
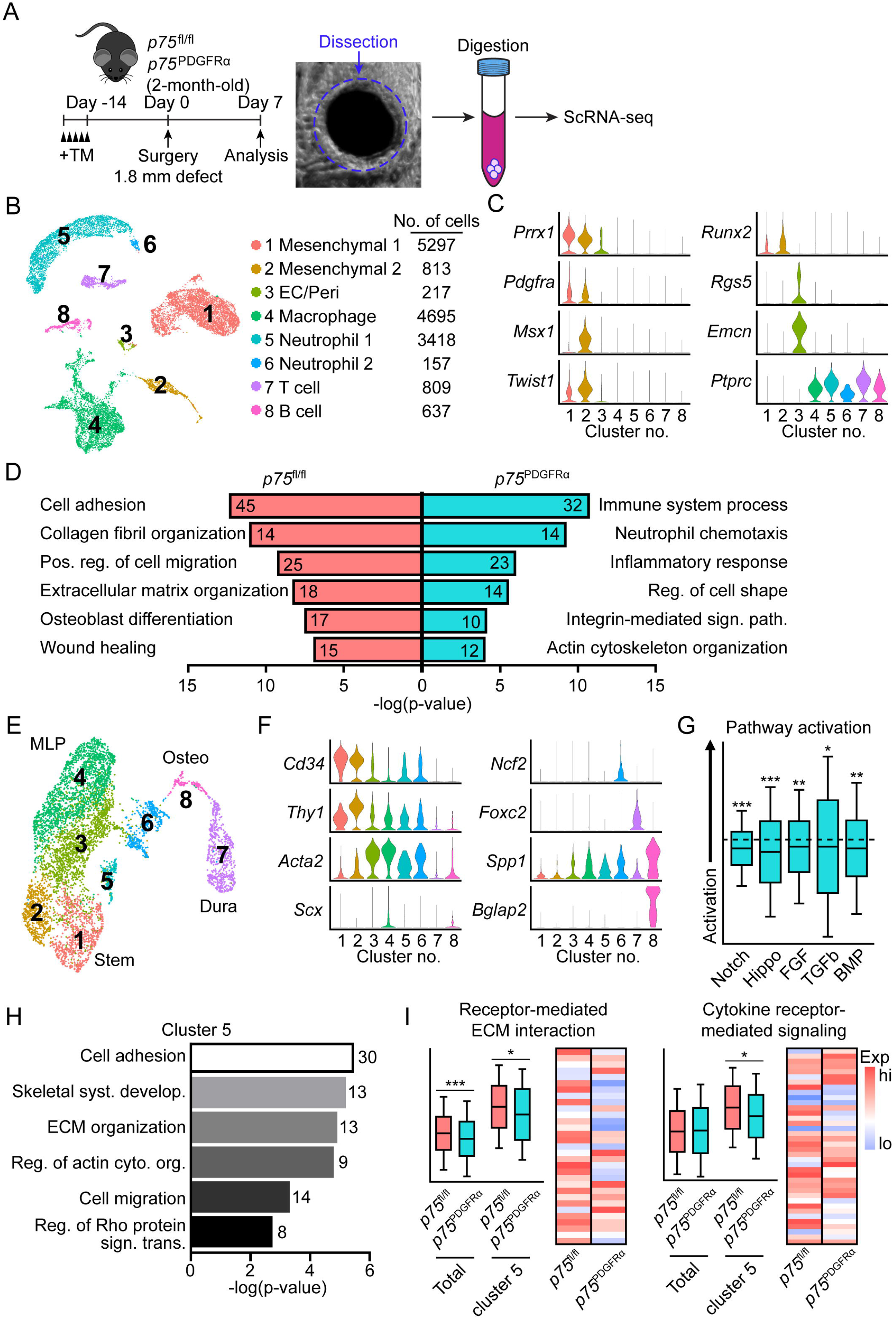
Single-cell transcriptomics highlights altered signaling pathway activation and impaired cellular migration with *p75* conditional gene deletion. (**A**) Schematic of experiment, in which *p75*^fl/fl^ or *p75*^PDGFRα^ animals (male, 2 mo old) were administered tamoxifen (TM), subjected to frontal bone injury after 14 d (circular calvarial defect, 1.8 mm diameter), with analysis performed 7 d post-operatively. The defect area was microdissected, dissociated and subjected to single-cell RNA sequencing. **(B)** UMAP plot of total cells isolated from calvarial defect site among *p75*^fl/fl^ and *p75*^PDGFRα^ animals. EC: endothelial cell; Peri: pericyte. **(C)** Violin plots of marker gene expression for mesenchymal, EC/Peri, and immune cell clusters. **(D)** GO term analyses of significantly enriched biological processes (up/downregulated GO terms) among all 16,043 cells from *p75*^fl/fl^ and *p75*^PDGFRα^ mice. The number in each column represents the number of enriched genes in the biological process. **(E)** UMAP plot of mesenchymal and neuroectodermal cells only (clusters 1 and 2 in subfigure B). MLP: Multilineage progenitor. **(F)** Violin plots of marker gene expression from mesenchymal subclusters. **(G)** Specific pathway analysis among mesenchymal cells from *p75*^PDGFRα^ mice in comparison to *p75*^fl/fl^ mice. Dashed black line indicates the same activation of pathway activity across *p75*^PDGFRα^ and *p75*^fl/fl^ genotypes. Normalized values reflect whether these pathways show positive (above dotted lines) or negative (below dotted lines) enrichment in *p75*^PDGFRα^ relative to *p75*^fl/fl^ samples. **(H)** GO term analyses of enriched biological processes among mesenchymal cluster 5. The number represents the number of enriched genes in the biological process. **(I)** Receptor-mediated ECM interaction and cytokine receptor-mediated signaling among total mesenchymal clusters or cluster 5 from *p75*^fl/fl^ and *p75*^PDGFRα^ mice. N=5 mice per group were used to yield a single-cell population. \**P*<0.05, \*\**P*<0.01, and \*\*\**P*<0.001 as assessed using a Wilcoxon test.

### p75 deletion inhibits cell migration and osteogenic differentiation in vitro

To determine the effect of p75 on stromal cell activity, primary neonatal mouse calvarial cells (NMCCs) from *p75*^fl/fl^ animals were isolated and treated with adenovirus encoding the Cre recombinase (Ad-Cre) or control (Ad-GFP). The efficiency of *p75* deletion was confirmed by qPCR (**Fig. 4A**) and validated by downstream signaling-related gene expression of RhoA, NF-κB, and MAPK signaling pathways (**Fig. S4**). Consistent with our *in vivo* analysis, NMCCs showed no change in cell proliferation rates among Ad-Cre- and Ad-GFP-treated groups (**Fig. 4B**). We did not observe an increase in apoptosis after Ad-Cre treatment by TUNEL assay (**Fig. 4C**). However, *p75* deletion reduced cell migration in wound-healing and transwell assays (**Fig. 4D,E**) as well as impaired osteogenic differentiation (**Fig. 4F,G**).

**Fig. 4.**
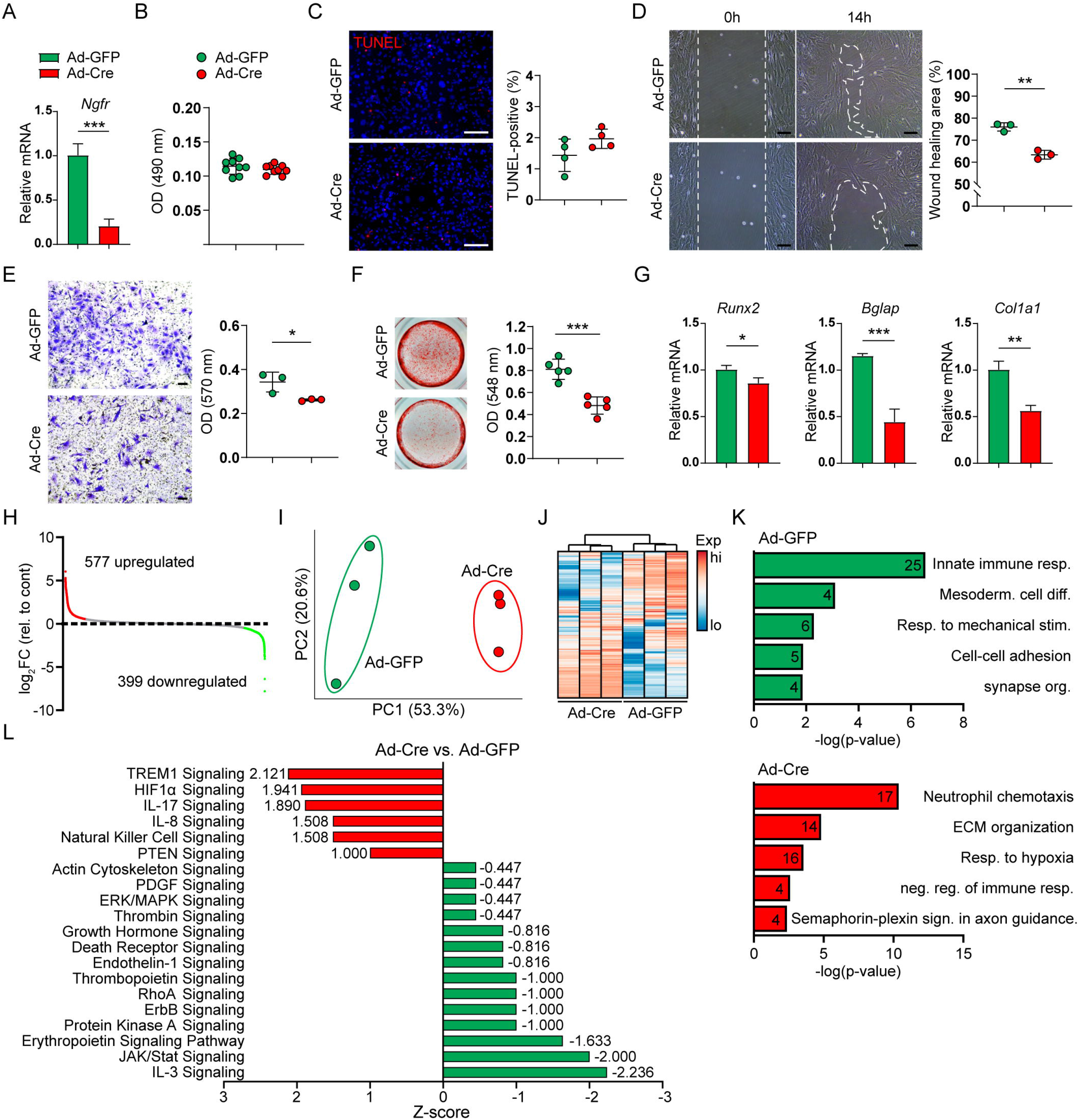
Deletion of *p75* in NMCCs inhibits cell migration and mineralization. NMCCs were isolated from *p75*^fl/fl^ mice and exposed to Adenoviral GFP (Ad-GFP) or Ad-Cre *in vitro*. **(A)** *Ngfr* expression by qRT-PCR among Ad-GFP- and Ad-Cre-treated cells after 72h. **(B,C)** Cellular proliferation by MTS assays (**B**, 72h) and apoptosis by TUNEL assay **(C)** among Ad-GFP- and Ad-Cre-treated NMCCs. **(D,E)** Cellular migration was assessed by **(D)** scratch wound healing assay at 14h or **(E)** transwell assay at 4h among Ad-GFP- and Ad-Cre-treated groups. For transwell assay, cells were placed on the membrane of the upper chamber, while medium was added to the bottom well. After 4h, cells were stained. **(F)** Osteogenic differentiation of Ad-GFP- and Ad-Cre-treated NMCCs by Alizarin Red staining and quantification at 14d. **(G)** Osteogenic gene expression among Ad-GFP- and Ad-Cre-treated NMCCs, including *Runx2*, *Bglap*, and *Col1a1* at 7d of differentiation. **(H-L)** Bulk total RNA sequencing among Ad-GFP- and Ad-Cre-treated NMCCs. **(H)** Differentially expressed genes (DEGs) of all 14,668 transcripts among Ad-Cre-treated NMCCs. Y-axis represents Log_2_fold change (FC). The number of upregulated DEGs (log_2_FC ≥ 1, red dots) is 577, the number of downregulated DEGs (log_2_FC ≤ −1, green dots) is 399. **(I,J)** Principal component analysis **(I)** and unsupervised hierarchical clustering **(J)** among Ad-GFP- and Ad-Cre-treated NMCCs. **(K)** DAVID functional GO analysis of biological processes enrichment among Ad-GFP- and Ad-Cre-treated NMCCs. The number in each column represents the number of enriched genes in the biological process. **(L)** QIAGEN Ingenuity Pathway Analysis identified representative pathways that were upregulated (Z-score > 0; red color) or downregulated (Z-score < 0; green color) in Ad-Cre compared to Ad-GFP-treated NMCCs. Dot plots represent an individual sample. Data are represented as mean ± 1SD. Scale bar: 100 μm (Black) and 10 μm (White). \**P*<0.05, \*\**P*<0.01, and \*\*\**P*<0.001 as assessed using a two-tailed Student’s *t*-test.

Next, the changes in the transcriptome of Ad-GFP- and Ad-Cre-treated NMCCs were examined using bulk RNA sequencing (RNA-seq). 14,668 protein-coding genes were expressed across all samples and had functional annotations. 577 transcripts, 3.93% of the total, showed a >2fold change (FC) increase among Ad-Cre-treated cells (red dots; 381 transcripts showed a significant increase (p<0.05)), while 399 transcripts, 2.72% of the total, showed a >2FC increase among Ad-GFP-treated cells (green dots; 219 transcripts showed a significant decrease (p<0.05)) (**Fig. 4H**). Quality control by principal component analysis (**Fig. 4I**) and hierarchical cluster of genes showing high sample deviation (**Fig. 4J**) confirmed strong reproducibility of sample replicas. Pathway analysis showed enrichment for GO terms linked to innate immune response, mesodermal cell differentiation, and cell-cell adhesion in Ad-GFP-treated cells, whereas neutrophil chemotaxis, extracellular matrix organization, and negative regulation of immune response were enriched in Ad-Cre-treated cells (**Fig. 4K**). Ingenuity Pathway Analysis (IPA) showed that the activated pathways in Ad-GFP-treated NMCCs are associated with the positive regulation of migration and osteogenesis, including IL-3 signaling, JAK/STAT signaling, and Erythropoietin signaling pathway (Z scores -2.236, -2, and -1.633; **Fig. 4L**) (*13–18*). Conversely, upregulated signaling pathways in Ad-Cre-treated NMCCs were associated with the negative regulation of cell migration and ossification, including IL-17 signaling as well as PTEN signaling (Z scores 1.89 and 1; **Fig. 4L**) (*19–21*). These data confirmed functional differences previously observed among *p75*^fl/fl^ and *p75*^PDGFRα^ mice, including the low migration and osteogenic potential of *p75*-deficient calvarial stromal cells.

### Inflammatory mesenchymal subpopulations are associated with shifts in macrophage phenotype with p75 conditional deletion

In addition to terms linked to migration and wound healing, pathway analysis of scRNA-seq and bulk RNA-seq revealed a recurring theme of altered immune processes in the context of *p75* deletion (**Fig. 5A**). To investigate how mesenchymal *p75* deletion may affect immune cell function, we examined the characteristics of mesenchymal subpopulations and found that mesenchymal subcluster 6 demonstrated DEGs enriched in several terms linked to immune cell recruitment and function (**Fig. 5B**, a full list of DEGs shown in **Table S3**). This mesenchymal subcluster 6 was found to express high levels of inflammatory regulators, such as *Il1a*, *Il10*, and *Tnf*, which were found to have reduced expression in cells derived from *p75*^PDGFRα^ mice (**Fig. 5C**).

**Fig. 5.**
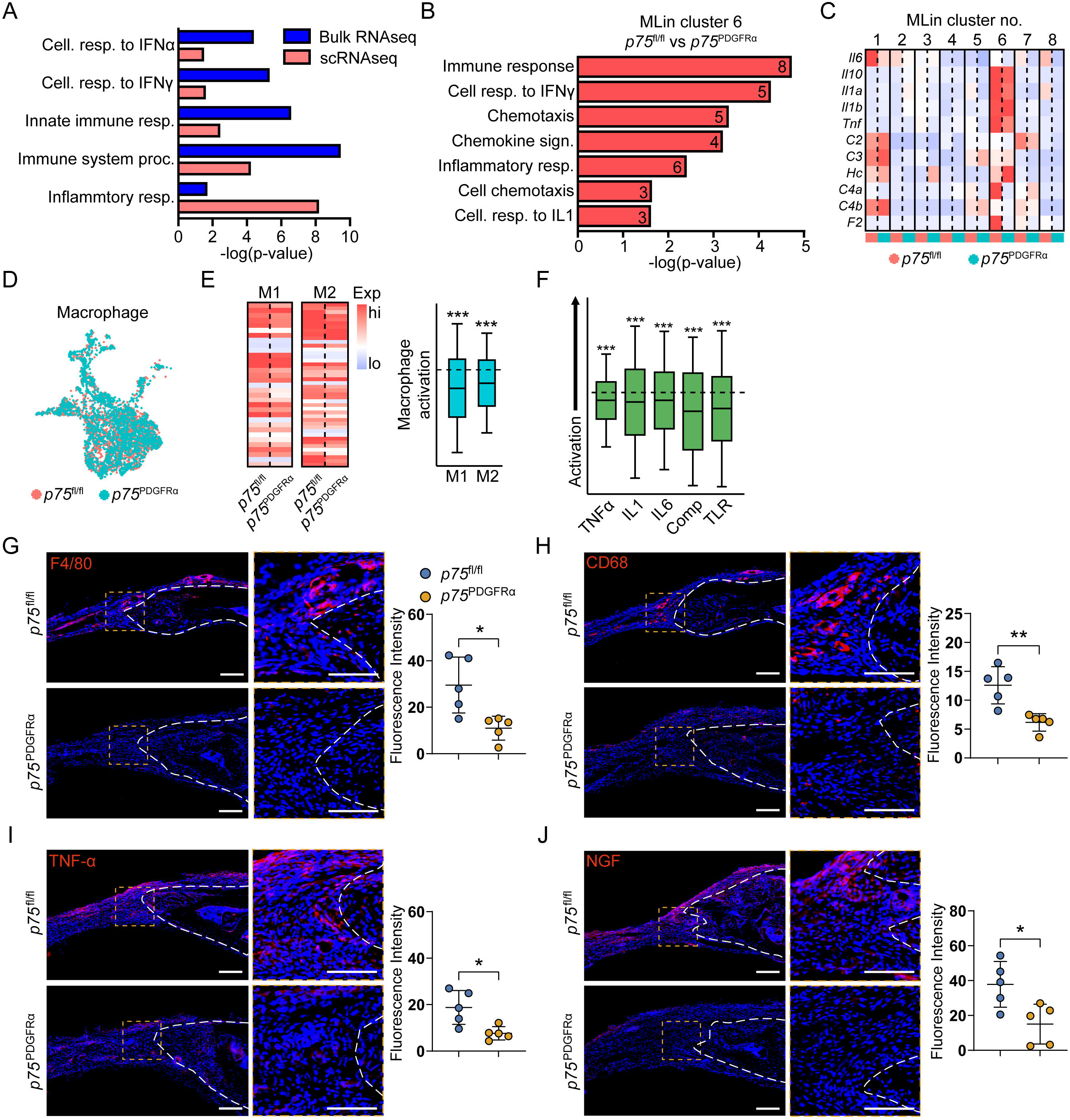
Deletion of p75 in PDGFRa-expressing mesenchyme results in secondary inflammatory changes in the bone defect microenvironment. **(A)** Inflammatory pathways enriched in both NMCC bulk RNA-seq and mesenchymal clusters derived from scRNA-seq. **(B)** GO term analyses of enriched biological processes among mesenchymal cluster 6. The number in each column represents the number of enriched genes in the biological process. **(C)** The expression of inflammatory factors among mesenchymal clusters from *p75*^fl/fl^ and *p75*^PDGFRα^ mice. **(D)** Distribution of macrophages from *p75*^fl/fl^ and *p75*^PDGFRα^ mice in UMAP plot. **(E)** Heat map of marker gene expression of M1 and M2 macrophages across *p75*^fl/fl^ and *p75*^PDGFRα^ mice (Left). The analysis of macrophage activation among *p75*^PDGFRα^ mice in comparison to *p75*^fl/fl^ mice. (Right). Dashed black line indicates the same activity of macrophage polarization across *p75*^fl/fl^ and *p75*^PDGFRα^ genotypes. **(F)** The expression of inflammatory factors among macrophages from *p75*^PDGFRα^ mice in comparison to *p75*^fl/fl^ mice. Normalized values reflect whether these pathways show positive (above dotted lines) or negative (below dotted lines) enrichment in *p75*^PDGFRα^ relative to *p75*^fl/fl^ samples. **(G-J)** Immunohistochemical staining and semi-quantitative analysis of F4/80 **(G)**, CD68 **(H)**, Tumor Necrosis Factor (TNF)-α **(I)**, and Nerve Growth Factor (NGF) **(J)** within the calvarial defect, d7 post-injury. Dashed white lines indicate bone edges. In graphs, each dot represents a single animal. White scale bar: 100 μm. N=5 mice per group for **(G-J)**. Data are represented as mean ± 1SD. \**P*<0.05, \*\**P*<0.01, and \*\*\**P*<0.001 as assessed using a Wilcoxon test or two-tailed Student’s *t*-test.

To determine whether a concomitant decrease was observed in macrophage signaling resulting from this reduced mesenchymal expression of inflammatory markers, we next investigated changes in macrophages within our scRNA sequencing of *p75*^fl/fl^ and *p75*^PDGFRα^ bone defects (**Fig. 5D-F**). While only a minor shift in overall population distribution was observed (**Fig. 5D, Fig. S2B**), expression of markers linked to both M1 and M2 macrophage activation were inhibited among *p75*^PDGFRα^ mice in comparison to *p75*^fl/fl^ mice (**Fig. 5E**). This was consistent with reductions in inflammatory pathways among macrophages within *p75*^PDGFRα^ defect sites, such as TNFα, IL1, IL6, the complement pathway, and Toll-like receptor signaling (**Fig. 5F**). In order to confirm this scRNA-seq data, macrophage number and activity were assessed by immunohistochemistry within *p75*^fl/fl^ and *p75*^PDGFRα^ bone defect sections. First, a clear reduction in F4/80^+^ (**Fig. 5G**) or CD68^+^ (**Fig. 5H**) macrophages was observed within *p75*^PDGFRα^ mice. TNF-α immunohistochemical staining was next performed, and confirmed a significant reduction across the *p75*^PDGFRα^ defect site (**Fig. 5I**). This diminished inflammatory setting was also associated with a reduction of NGF immunoreactive cells with the *p75*^PDGFRα^ defect site (**Fig. 5J**). In aggregate, depletion of *p75* in PDGFRα-expressing mesenchymal cells significantly inhibits injury-associated macrophage activation and NGF expression, associated with a delay in bone defect repair.

### p75 promotes human calvarial osteoblasts migration and osteogenesis

Finally, the role of p75 in human cells and tissue samples was examined (**Fig. 6, Table S4**). To investigate the role of p75 in human calvarial osteoblasts, siRNA-mediated knockdown of *p75* was performed (**Fig. 6A**). As in mouse NMCCs, knockdown of *p75* inhibited cell migration in wound-healing and transwell assay invasion (**Fig. 6B,C**) as well as impaired osteogenic differentiation (**Fig. 6D,E**). Next, detection of p75 expression was observed across human fracture calluses (**Fig. 6F**, N=3 human fracture calluses, see demographic information in **Table S4**). In similarity to findings in mouse calvariae, p75 was abundantly expressed in bone lining cells of the fracture callus, especially in Osteocalcin^+^ bone lining osteoblasts (**Fig. 6G,H**). Thus, *p75* in osteoblasts and their cellular precursors is essential for cell migration and bone repair.

**Fig. 6.**
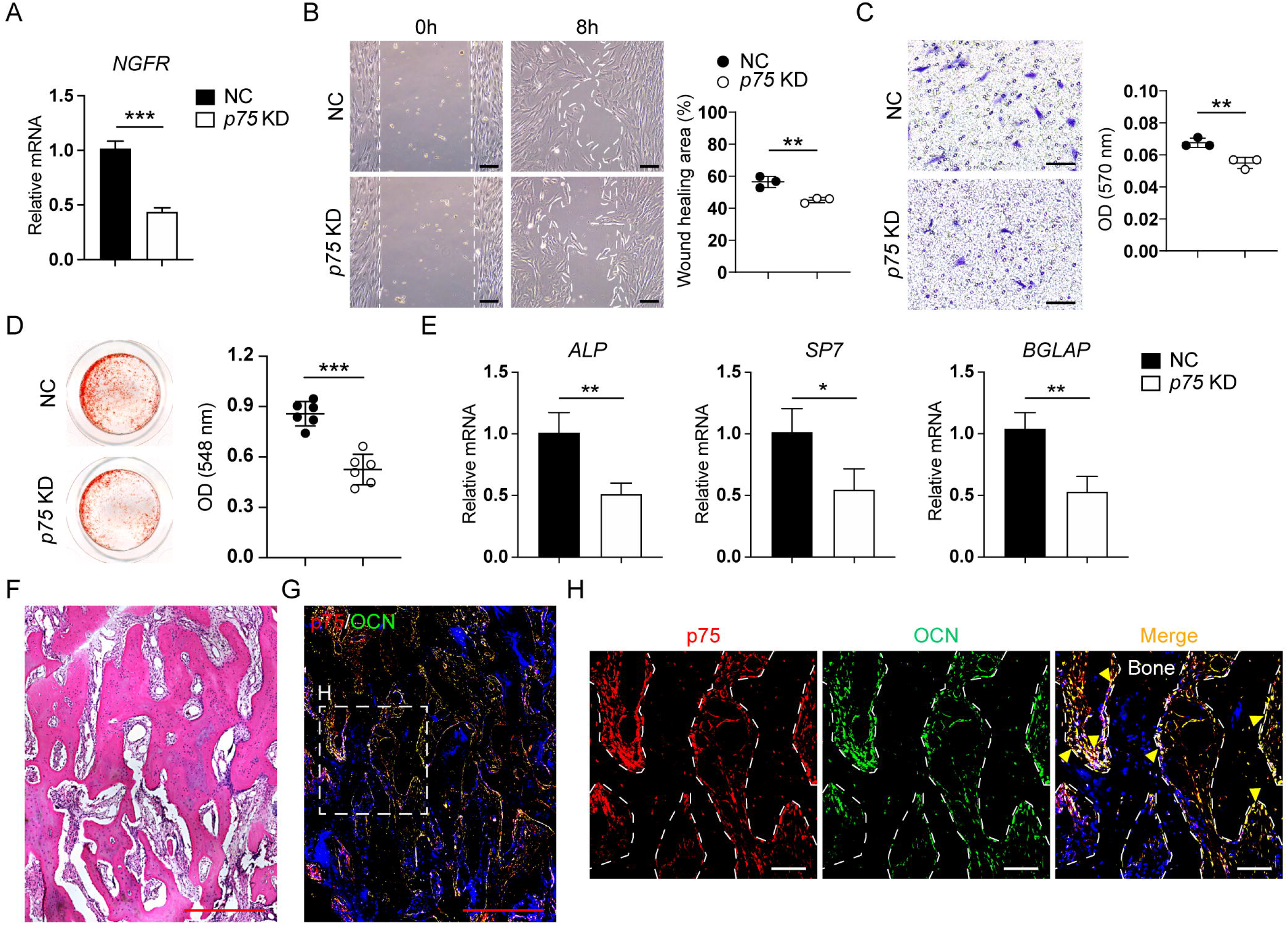
Deletion of *p75* in human calvarial osteoblasts inhibits cell migration and mineralization. **(A)** *NGFR* expression by qRT-PCR after siRNA-mediated knockdown in human calvarial osteoblasts. NC, negative control (non-targeting siRNA). **(B,C)** Cellular migration was assessed by **(B)** scratch wound healing assay at 8 h or **(C)** transwell assay at 4 h with or without *p75* siRNA. Representative 100x images with percentage gap closure are shown. **(D)** Osteogenic differentiation after siRNA-mediated knockdown in human calvarial osteoblasts by Alizarin Red (AR) staining and quantification at 14 d. **(E)** Osteogenic gene expression with or without *p75* siRNA, including *Alkaline phosphatase* (*ALP*), *Osterix* (*SP7*), and *Osteocalcin* (*BGLAP*) at 7 d of differentiation. **(F)** Representative histologic appearance of a healing human fracture by H&E staining. **(G)** Immunohistochemistry for p75 (red) and Osteocalcin (OCN, green) in healing human fracture sites (N=3). Nuclei, DAPI (blue). Dashed box indicates the location of H. **(H)** High magnification image for p75 (red) and OCN (green) expression in the bone lining osteoblasts. Yellow arrowheads indicate p75 and OCN co-localization. White dashed lines indicate bone edges. Graphs represent mean values, while error bars represent 1 SD. Dot plots represent an individual sample, while whisker plots indicate mean values and 1SD. *In vitro* experiments were performed in biologic and experimental triplicate. Red scale bar: 500 μm. Black/white scale bar: 100 μm. \**P*<0.05, \*\**P*<0.01, and \*\*\**P*<0.001 as assessed using a two-tailed Student’s *t*-test.

## Discussion

In this study, we sought to define the cellular and molecular events that accompany bone repair in genetic mouse models that enabled specific disruption of p75 signaling components in bone. NGF is acutely upregulated in the early period following bone injury (*3*), triggering a migration of p75-expressing mesenchymal cells into the damaged area. When this signaling pathway is disrupted, defective cellular migration, ossification, and a secondary blunting of the immune response with changes in macrophage activation are the result. Given these observations, it is not clear why experimental work with neutralizing antibodies to NGF has not shown a significant delay in bone healing (*22, 23*). It is probable that the analgesic effects of anti-NGF are observed at a lower dose than those needed to impede tissue repair. Nevertheless, when taken together with our past work, these data illustrate the multimodal and critical effects of NGF in bone repair, including coordinating proper mesenchymal cell migration, re-innervation and vascularization, and ossification – cellular functions which are regulated in concert via both NGF receptors.

Our studies focused on the role of p75 in mesenchymal progenitors and osteoblasts by using available transgenic animals. However, p75 expression is of course present in other cell types, including pericytes and immune cells. For example, proNGF-p75 signaling in the vasculature, including pericytes and endothelial cells, has been shown to mediate vascular response to injury of either the myocardium or injured limb in ischemia-reperfusion models (*24–26*). Our observations on unchanged vascularity in the context of *p75* conditional gene deletion thus likely stem from the choice of Cre driver. Likewise, macrophages are also responsive to the chemotactic properties of proNGF via the low-affinity receptor p75 (*27*), and demonstrate an altered secretome with minor changes in M1/M2 phenotype after proNGF treatment (*27*). Thus, it is possible that p75 has more functional roles in tissue repair than would be illustrated by our experimental design. Second, our studies were focused on a single neurotrophin given our laboratories’ interests and past work in this area (*2–4, 28*). Nevertheless, other neurotrophins such as BDNF (Brain-derived neurotrophic factor) are also expressed in human fracture sites including in endothelial and osteoblastic cells during early repair phases (*29*). Indeed, our sequencing analysis identified a host of transcripts for neurotrophins, neuroattractants and neurorepellents present within the calvarial injury site, such as Netrins and Semaphorins. It is likely that several neurotrophins or other axonal guidance molecules have functionally redundant roles in regulating tissue ingrowth after bone injury – a hypothesis that has not yet been analyzed experimentally.

In this study, we computationally identified several subpopulations of mesenchymal calvarial cells within a bone defect site. Some subclusters showed characteristic genes associated with a stage of mesenchymal cell differentiation, such as enrichment for stem/progenitor markers like *Thy1*, or conversely mature osteoblast markers, such as *Bglap2*. Two unique cell subclusters were also identified which were further shown to have significant transcriptional changes within the context of *p75* conditional deletion. Here, subcluster 5 appeared to be enriched in ‘migrating’ mesenchymal cells based on specific enrichment in GO terms related to interaction between cells and their surrounding matrix, as well as increased responsiveness to chemokine-mediated signaling. Likewise, subcluster 6 was defined as an ‘immunomodulatory’ cluster based on the enriched expression of inflammatory cytokines such as *Il10, Il1a/b* and *Tnfα*. This cluster may have similarities to fibroblastic progenitor cells previously identified in dermal wounds, which demonstrate an inflammatory gene signature by single-cell analysis (*30*). Further work is required to determine whether these mesenchymal subclusters indeed represent unique cell types, or instead simply reflect a unique functional response to tissue injury.

The extent to which NGF signaling can be targeted therapeutically for its beneficial effects on bone and bone repair represents an evolving area of interest in the orthopaedic research community. Several recent papers suggest that positive regulation of NGF-TrkA signaling may have beneficial effects in long bone fracture repair. For example, local injection of NGF stimulated early fracture maturation in a closed tibial fracture mouse model (*31*). Systemic administration of the partial TrkA agonist Gambogic amide has been used to improve indices of fibular fracture healing in mice (*32*), and analogously to augment skeletal adaptation to mechanical load (*33*). Our present study demonstrates the multiple cell-specific functions that NGF signaling plays in bone repair. In combination with past work (*2, 3*), we document clear and potentially synergistic roles for NGF-p75 and NGF-TrkA signaling in osseous repair.

## Materials and Methods

### Mice

All animal experiments were performed according to approved protocols (MO16M226 & MO19M366) of the Animal Care and Use Committee (ACUC) at Johns Hopkins University (JHU). Pdgfrα-CreER^TM^ animals were a kind gift from the Dwight Bergles laboratory (*34*) and are commercially available (The Jackson Laboratory, Stock No. 018280, Bar Harbor, ME). Pdgfrα^mT/mG^ mice were obtained by crossing Pdgfrα-CreER^TM^ with mT/mG mice (JAX Stock No. 007576). *p75*^PDGFRα^ mice were obtained by crossing Pdgfrα^mT/mG^ mice with *p75*^fl/fl^ mice (JAX Stock No. 031162). Tamoxifen (TM; Sigma-Aldrich, St. Louis, MO) was dissolved in sunflower seed oil (Sigma-Aldrich) and injected intraperitoneally according to previously validated protocols (TM: 150 mg/kg/d for 5 d) (*12, 35*). LysM-Cre mice were purchased from the Jackson Laboratory (JAX Stock No. 004781). In order to achieve cell-specific deletion of *Ngf*, *Ngf*^LysM^ were obtained by crossing LysM-Cre animals with *Ngf*^fl/fl^ animals. When feasible, littermate analysis was performed by investigators blinded to the mouse genotype. All these lines were backcrossed for at least 8 generations to the C57BL/6J background before used for the experiments reported here.

### Calvarial defect procedures

Calvarial defects were performed based on our prior methods (*3, 35, 36*). In Pdgfrα^mT/mG^ or *p75*^PDGFRα^ mice, TM administration was performed 14 d prior to defect creation as previously validated (*12, 35*). Mice were provided isoflurane (2-3% inh) and Buprenorphine (1 mg/kg sc). Briefly, hair overlying the calvaria was clipped, and a 1 cm skin incision was made over the midline skull to expose the frontal bone. Next, a 1.8 mm diameter, full-thickness, circular frontal bone defect was created in the non-suture associated frontal bone (right side) using a micro surgical drill and a trephine drill bit. Meticulous care was taken to protect the neighboring sutures and the underlying dura mater. Finally, the skin was sutured and the animals were monitored per established postoperative protocols. Mice were euthanized after 3, 7 or 28 d for analysis.

### Radiographic analyses

Skulls were fixed in 4% paraformaldehyde (PFA) for 24 h and evaluated using a high-resolution micro-CT imaging system (SkyScan 1275; Bruker, Kontich, Belgium). Scans were obtained at an image resolution of 12 μm and set as 1 mm of aluminum filter, X-ray voltage of 65 kVp, anode current of 153 μA, exposure time of 160-218 ms, frame averaging of 6, and rotation step of 0.3 degrees. Three-dimensional images were then reconstructed from the 2D X-ray projections using a commercial software NRecon software (v1.7.0.4, SkyScan). For 3D morphometric analyses of images, CTAn software (v1.16, SkyScan), CTVol (v2.0, SkyScan), and CTVox (v3.2, SkyScan) were used. For calvarial defect analysis, a cylindrical volume of interest centered around each defect site was defined as the 1.8 mm in diameter and 1.2 mm in height with a threshold value of 70-255. Bone volume (BV) and fractional bone volume (bone volume/tissue volume (BV/TV)) were calculated from binary x-ray images. Lastly, bone fractional area (BFA) and defect diameter were calculated by using CTVox to create a 3D rendering of calvarial defect and measuring by ImageJ software (Version 1.8.0; NIH, Bethesda, MD).

### Histology and immunohistochemistry

After radiographic imaging, samples were decalcified in 14% EDTA for 21 days, embedded in optimal cutting temperature compound (OCT; Sakura, Torrance, CA), and sectioned in a coronal plane at 20 or 50 um thickness. H&E staining was performed on serial sections. For immunofluorescent staining, all sections were incubated with trypsin enzymatic antigen retrieval solution (Abcam, Cambridge, MA, USA) for 3 min at room temperature (RT). When 50 µm sections were used, sections were next permeabilized with 0.5% Triton-X for 30 min. All sections were blocked with 5% goat or donkey serum in PBS for 1 h at RT and incubated with the primary antibodies (see **Table S5** for a summary of antibodies used) at 37 °C for 3 h or overnight at 4 °C. Next, secondary antibodies (1:200) were used with incubation for 2 h at RT. Sections were counterstained with DAPI mounting medium (Vector Laboratories, Burlingame, CA). All histological sections were examined under a Zeiss 800 confocal microscope (Zeiss, Thornwood, NY) or Leica DM6 microscope (Leica Microsystems Inc, Wetzlar, Germany). Staining without primary antibody was used to exclude non-specific staining. For visualization of reporter activity, a wildtype specimen prepared in the same fashion was used as a control to identify the intensity of autofluorescence. Images were quantified using ImageJ or Imaris (Bitplane, Belfast, UK) software. For the lineage tracing of cell migration and apoptosis, the percentage of PDGFRα^+^ cells was calculated in the central defect region. For the TUBB3 staining, the volume of TUBB3^+^ nerve fibers was quantified with three different sites of the defect span and the average value was taken as the final value of the sample. For all other immunohistochemistry, the fluorescence intensity was quantified on the edges of bone in single microscopical fields.

### Isolation and culture of neonatal mouse calvarial cells

Primary neonatal mouse calvarial cells (NMCCs) were collected from *p75*^fl/fl^ or *p75*^PDGFRα^ embryos at postnatal day 2. Frontal and parietal bones were minced and subjected to six sequential enzymatic digestions with a mixture containing 1 mg/mL collagenase type I (Worthington Biochemical Corporation, Lakewood, NJ; LS004197) and 1 mg/mL collagenase type II (Worthington Biochemical Corporation; LS004177) (*37*). Cell fractions (from sequential digestions 3-6) were collected and cultured in α-MEM supplemented with 15% (vol/vol) FBS, 100 U/mL penicillin, and 100 mg/mL streptomycin. In select experiments, *p75*^fl/fl^ NMCCs were exposed to an adenovirus encoding either Cre Recombinase (Ad-Cre, multiplicity of infection=150; 1045-HT, Vector Biosystems, Malvern, PA) or GFP (Ad-GFP, multiplicity of infection=150; 1060-HT, Vector Biosystems). In additional select experiments, *p75*^PDGFRα^ NMCCs were treated with 4-Hydroxytamoxifen (2 μM) or vehicle control (Sigma-Aldrich).

### Proliferation

Proliferation assays were performed in 96 well plates (2 × 10^3^ NMCCs/well) and assayed at 48 h using the CellTiter96® AQueous One Solution Cell Proliferation Assay kit (MTS; Promega, Madison, WI) (*38*). Briefly, 20 μl of MTS solution was added to each well. After incubation for 1 h at 37 °C, the plate was measured using an Epoch microspectrophotometer (BioTek, Winooski, VT) by absorbance at 490 nm.

### Isolation of mouse macrophages and macrophage conditioned medium

Macrophages were collected from *Ngf*^fl/fl^ or *Ngf*^LysM^ mice (8 weeks old mixed-gender animals). Bone marrow cells were flushed from the long bones (femur and tibia) with culture medium and resuspended in medium containing macrophage colony-stimulating factor (M-CSF; 15 ng/mL) (*39*). After 7 d in culture, adherent bone marrow-derived macrophages were incubated in growth medium without M-CSF for 2 d and the supernatant was collected for conditioned medium experiments.

### Migration

Cell migration was measured with Ibidi inserts (Ibidi, Planegg/Martinsried, Germany), where 2 × 10^4^ NMCCs or human calvarial osteoblasts (purchased from ScienCell Research Laboratories, Carlsbad, CA; Catalog #4600) were seeded and grown to confluency. SiRNA-mediated knockdown of *NGFR* was performed among human calvarial osteoblasts. Human *NGFR* siRNA or negative control siRNA were obtained from ThermoFisher Scientific (Waltham, MA). SiRNA was transfected using TransIT®-LT1 Transfection Reagent (Mirus Bio, Madison, WI) as described by the manufacturer. Inserts were removed and cell migration into the empty area was monitored by brightfield microscopy at 0, 8, or 14 h. The equilibrium width of the gap was calculated using the ImageJ software. Here, gap closure = (scratch area at hour 0 − scratch area at hr 8 or 14)/scratch area at hr 0 × 100%. For the transwell migration assays, 2.0 × 10^4^ NMCCs or human calvarial osteoblasts were resuspended in 100 μL of α-MEM medium without FBS and were placed in the upper well of 8-μm pore-size Transwell 24-insert plates (Corning, NY). The migration assay was performed by adding to the bottom well 600 μL of α-MEM medium supplemented with 15% FBS or conditioned medium from cultured macrophages. After 4 h, cells on the bottom of the inserts were fixed and stained with 0.5% crystal violet (Sigma-Aldrich) for 15 min. Pictures were taken by using 100× magnification (Leica Microsystems Inc). Crystal violet was eluted using 33% acetic acid, and the eluent was quantified by absorbance at 570 nm.

### ELISA assay

Macrophages derived from *Ngf*^fl/fl^ or *Ngf*^LysM^ mice were cultured with M-CSF for 7 d. Then, adherent bone marrow-derived macrophages were incubated in the culture medium with 1% FBS for 2 d and the cell lysates were collected for ELISA assay. The concentrations of NGF were detected by ELISA kit (Biosensis Pty Ltd, Thebarton, South Australia) according to the manufacturer’s protocol.

### Osteogenic differentiation

NMCCs or human calvarial osteoblasts were seeded in 24-well plates at a density of 1 × 10^5^ cells/well. Osteogenic differentiation medium (ODM) consisted of α-MEM, 10% FBS, 1% penicillin/streptomycin with 100 nM dexamethasone, 50 μM ascorbic acid, and 10 mM β-glycerophosphate (*40–42*). 24 h after cell seeding, growth medium was replaced with ODM, replenished every 3-4 d. For alizarin red (AR) staining, cells were fixed with 4% paraformaldehyde and stained with a 2% alizarin red solution at 14 d of differentiation. Pictures were taken using Olympus Epson scanner (Los Angeles, CA), followed by incubation with 0.1 N sodium hydroxide and quantified using an Epoch microspectrophotometer (BioTek) by absorbance at 548 nm.

### Ribonucleic acid isolation and quantitative real-time polymerase chain reaction

Total RNA was extracted from the cultured cells using TRIzol Reagent (Invitrogen, Carlsbad, CA, USA) according to the manufacturer’s instructions. 1 μg of total RNA was used for reverse transcription with iScript™ cDNA synthesis kit (Bio-Rad, Hercules, CA) following manufacturer’s instructions. Real-time PCR was performed using SYBR™ Green PCR Master Mix (ThermoFisher Scientific) according to the manufacturer’s protocol. Relative gene expression was calculated using a 2^-ΔΔCt^ method by normalization with *GAPDH*. Primer sequences are presented in **Table S6**.

### Transcriptomics

The RNA content of Ad-Cre and Ad-GFP treated NMCCs was detected by total RNA sequencing. Briefly, *p75*^fl/fl^ NMCCs were exposed to Ad-Cre or Ad-GFP (multiplicity of infection=150) for 72h, then total RNA was extracted from Ad-Cre and Ad-GFP treated *p75*^fl/fl^ NMCCs by Trizol (Life technologies corporation, Gaithersburg, MD). The RNA samples were sent to the JHMI Transcriptomics and Deep Sequencing Core and quantified by deep sequencing with the Illumina NextSeq 500 platform (Illumina, San Diego, CA). Data analyses were performed using software packages including Partek Genomics Suite, Spotfire DecisionSite for Functional Genomics, QIAGEN Ingenuity® Pathway Analysis, and DAVID bioinformatics resources (*43*).

### Apoptosis detection

Ad-Cre and Ad-GFP treated *p75*^fl/fl^ NMCCs were seeded in Millicell EZ SLIDE (MilliporeSigma, Burlington, MA) and cultured for 24 h. The cells were fixed in 4% PFA for 15 min and permeabilized with 0.25% Triton X-100 for 20 min. TUNEL assay was carried out using the Click-iT Plus TUNEL Assay for in situ apoptosis detection with Alexa Fluor 647 dyes (ThermoFisher Scientific). Pictures were taken using a Leica DM6 microscope.

### Flow cytometry analysis

In order to detect cell migration *in vivo*, skull defects were microdissected 7 days after injury. The cells in the defect site were obtained by collagenase Type I/II (1 mg/mL) digestion using the above method. Briefly, cell fractions (from sequential digestions 1-6) were collected and resuspended in red blood cell lysis buffer (37°C for 5 min). After centrifugation, cells were resuspended in Hanks’ Balanced Salt Solution (HBSS) and 0.5% bovine serum albumin. The resulting cells were processed for cell sorting, using a mixture of the following directly conjugated antibodies (**Table S5**): anti-CD31-allophycocyanin (1:30), anti-CD45-allophycocyanin (1:30), anti-Ter119-allophycocyanin (1:30), anti-PDGFRα-BV421 (1:50) were added separately and incubated at 4°C for 20 min. Cells were then washed with PBS and examined with a DakoCytomation MoFlo (Beckman, Indianapolis, IN). FlowJo software (Tree Star Inc., Ashland, OR) was used for the analysis of flow cytometry data. Gating and PDGFRα frequency were set in relation to matching isotype control.

### Single-cell RNA sequencing

Skulls were microdissected 7 days after defect creation and digested with collagenase Type I/II (1 mg/mL) digestion using the above method. Cell fractions (from sequential digestions 1-6) were collected and resuspended in red blood cell lysis buffer (37°C for 5 min). Digestions were subsequently filtered through 40μm sterile strainers. Cells were then washed in PBS and resuspended in HBSS at a concentration of ∼1000 cells/μl. Cell viability was assessed with Trypan blue exclusion on a Countess II (ThermoFisher Scientific) automated counter and showed a >85% viability. Cells were sent to the JHMI Transcriptomics and Deep Sequencing Core. The library was generated using the 10X Genomics Chromium controller following the manufacturer’s protocol. Cell suspensions were loaded onto a Chromium Single-Cell A chip along with reverse transcription (RT) master mix and single-cell 3’ gel beads, aiming for 10,000 cells per channel. Following generation of single-cell gel bead-in-emulsions (GEMs), reverse transcription was performed and the resulting Post GEM-RT product was cleaned up using DynaBeads MyOne Silane beads. Beads are provided by 10X with the kit. The cDNA was amplified, SPRIselect (Beckman Coulter, Brea, CA) cleaned and quantified then enzymatically fragmented and size selected using SPRIselect beads to optimize the cDNA amplicon size prior to library construction. An additional round of double-sided SPRI bead cleanup is performed after end repair and A-tailing. Fragmentation, end repair and a-tailing are all one reaction, followed by the double-sided cleanup. Another single-sided cleanup is done after adapter ligation. Indexes were added during PCR amplification and a final double-sided SPRI cleanup was performed. Libraries were quantified by Kapa qPCR for Illumina Adapters (Roche) and size was determined by Agilent Bioanalyzer 2100. Read 1 primer, read 2 primer, P5, P7, and sample indices were incorporated per standard GEM generation and library construction via end repair, A-tailing, adaptor ligation and PCR. Libraries were generated with unique sample indices (SI) for each sample. Libraries were sequenced on an Illumina NovaSeq SP 100 cycle (San Diego, CA). *CellRanger* was used to perform sample de-multiplexing, barcode processing, and single-cell gene counting (Alignment, Barcoding, and UMI Count) at the JHMI Transcriptomics and Deep Sequencing Core. Downstream analysis steps were performed using *Seurat*. Cells were first filtered to have >500 and <8,000 detected genes, as well as less than 20% mitochondrial transcripts. *SCTransform*, including regression for cell cycle scores derived using the *CellCycleScoring* function, and dimensional reductions using uniform manifold approximation and projection (UMAP) was performed using *Seurat*. Pathway activation or module scores were generated using the *AddModuleScore* function of *Seurat* using validated gene lists from KEGG pathways. Module scores were calculated as the level of gene expression enrichment of a set gene list relative to a random control list, with higher module score values representing positive enrichment beyond background. Specific gene lists for module scores are provided in **Tables S7 and S8**.

### Statistical analysis

Data are presented as the mean ± one standard deviation (SD). Statistical analyses were performed using GraphPad Prism (Version 7.0) or R package *ggpubr*. *In vitro* experiments were performed in biologic and experimental triplicate. The number of animals used in the *in vivo* experiments is shown in figure legends. For cell depletion experiments, our depletion efficiency resulted in effect sizes of 1.75 or higher. For these scenarios, with at least 4 mice per group, a two-sample *t*-test would provide 80% power to detect effect sizes of at least 1.75, assuming a two-sided 0.05 level of significance. The Kolmogorov–Smirnov test was used to confirm normal distribution of the data. A two-tailed Student’s *t*-test or Wilcoxon test was used for two-group comparisons. A one-way ANOVA test was used for multiple groups, followed by Tukey’s multiple comparisons test. \**P*<0.05, \*\**P*<0.01, and \*\*\**P*<0.001 were considered significant.

## Acknowledgments

We thank the JHU microscopy core facility, JHMI deep sequencing and microarray core facility, and Hao Zhang within the JHU Bloomberg Flow Cytometry and Immunology Core.

## Author contributions

Conceptualization: JX, ZL, RJT, AWJ

Methodology: JX, ZL, SN, YW, CAM, TS, QQ, XX

Investigation: EFM

Visualization: RJT, AL

Supervision: TLC, AWJ

Writing—original draft: JX, RJT, AWJ

Writing—review & editing: TLC, AWJ

## Data and materials availability

Expression data that support the findings of this study have been deposited in Gene Expression Omnibus (GEO) with the accession code GSE179891. All data needed to evaluate the conclusions in the paper are present in the paper and/or the Supplementary Materials.

