## Supplementary Figures and Tables for "NGF-p75 signaling coordinates skeletal cell migration during bone repair"

**This PDF file includes:**

Figs. S1 to S4
Tables S1 to S8

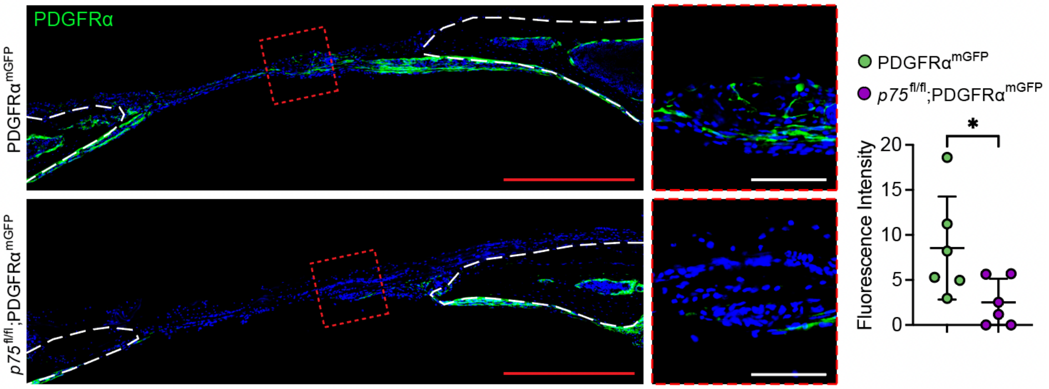

**Fig. S1. Mesenchymal cell migration, as assessed by Pdgfrα-CreER; mT/mG cell lineage tracing.** Tile scan (left) and high-magnification images of the central defect (right) demonstrate the migration of GFP$^{+}$ (PDGFRα$^{mGFP}$) progenitor cells into the defect site at d3 post-injury. GFP intensity was quantified in the defect. Dashed white lines indicate bone edge. Tdtomato is not shown. In graphs, each dot represents a single animal. Red scale bar: 500 µm. White scale bar: 100 µm. N=6 animals per group. Data are represented as mean ± 1 SD. *$P<0.05$ as assessed using a two-tailed Student's *t*-test.

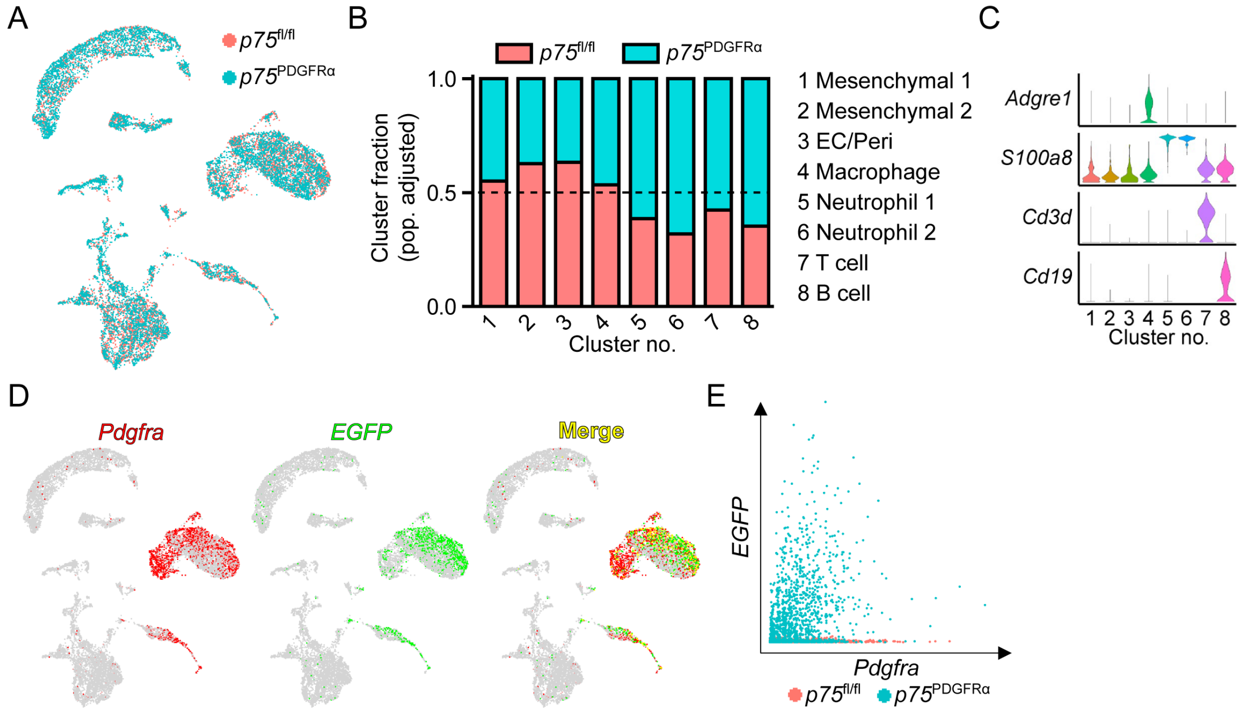

**Fig. S2. Additional description of cell identity and frequency by single-cell RNA sequencing, related to Figure 3.** Seven days after frontal bone defect creation, the injury site was microdissected and cells were dissociated and subjected to scRNA-seq. **(A)** Distribution of total cells derived from $p75^{fl/fl}$ (red) and $p75^{PDGFR\alpha}$ (blue) mouse calvarial defects as shown by UMAP plot. **(B)** The normalized frequency of cells from $p75^{fl/fl}$ and $p75^{PDGFR\alpha}$ mice in each cluster. The dashed line indicates an even distribution of cells across $p75^{fl/fl}$ and $p75^{PDGFR\alpha}$ genotypes. **(C)** Violin plots of marker gene expression for macrophage, neutrophil and lymphocyte cell clusters. **(D)** Validation of Cre-recombination in Pdgfr$\alpha^{+}$ cells. UMAP projections demonstrating expression of *Pdgfra* in clusters 1 and 2, overlapping with *eGFP* transcripts in clusters 1 and 2. **(E)** Scatter plot showed that EGFP was only expressed in cells from $p75^{PDGFR\alpha}$ (blue) mice. N=5 mice per group were used to yield a single cell population. Analyses were among all 16,043 cells from $p75^{fl/fl}$ and $p75^{PDGFR\alpha}$ mice.

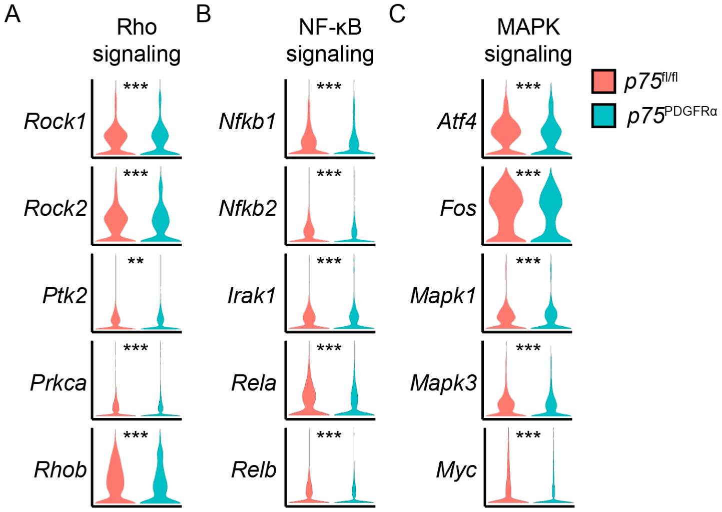

**Fig. S3. Expression of p75 downstream signaling pathways in mesenchymal cells among *p75*$^{fl/fl}$ and *p75*$^{PDGFR\alpha}$ mice, related to Figure 3.** Violin plots shown derived from scRNA sequencing of microdissected calvarial defects. **(A)** The expression of Rho signaling-related genes, including *Rho-associated coiled-coil containing protein kinase 1* (*Rock1*), *Rock2*, *Protein tyrosine kinase 2* (*Ptk2*), *Protein kinase C alpha* (*Prkca*), and *Ras homolog family member b* (*Rhob*). **(B)** The expression of Nuclear factor-kappa B (NF-κB) signaling-related genes, including *NF-κB subunit 1* (*Nfkb1*), *Nfkb2*, *Interleukin 1 receptor associated kinase 1* (*Irak1*), *Rela*, and *Relb*. **(C)** The expression of MAPK signaling-related genes, including *Activating transcription factor 4* (*Atf4*), *Fos*, *Mitogen-activated protein kinase 1* (*Mapk1*), *Mapk3*, and *Myc*. **$P<0.01$ and ***$P<0.001$ as assessed in R package *ggpubr* using a Wilcoxon test.

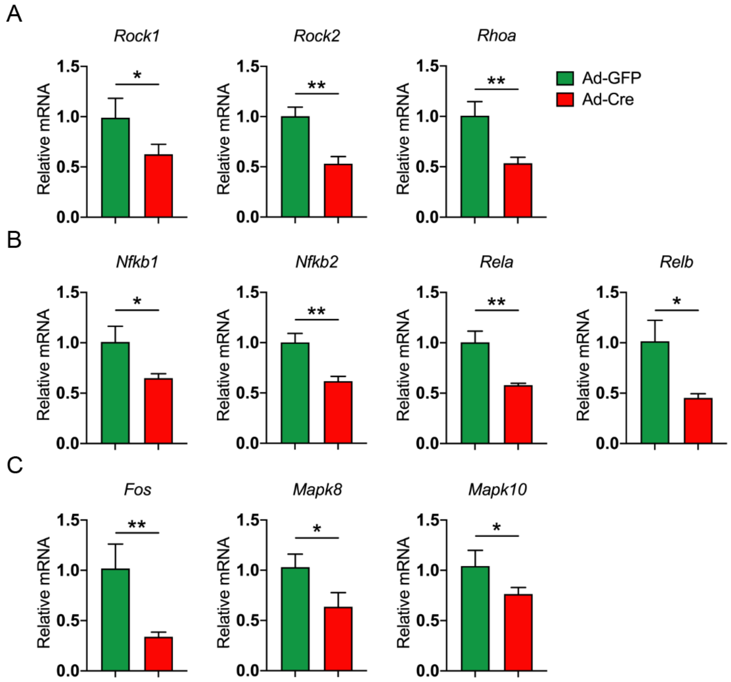

**Fig. S4. The expression of p75 downstream-related genes among Ad-GFP- and Ad-Cre-treated NMCCs, related to Figure 4. (A)** The expression of Rho signaling-related genes, including *Rock1*, *Rock2*, and *Rhoa*, by qRT-PCR. **(B)** The expression of NF-κB signaling-related genes, including *Nfkb1*, *Nfkb2*, *Rela*, and *Relb*. **(C)** The expression of MAPK signaling-related genes, including *Fos*, *Mapk8*, and *Mapk10*. *$P$<0.05 and **$P$<0.01 as assessed using a two-tailed Student's *t*-test.

**Table S1.** The frequency of lineage-negative ($Lin^-$) and $Lin^+$ among freshly isolated calvarial tissue in *Ngf*$^{fl/fl}$ and *Ngf*$^{LysM}$ mice.

| | Cell frequency among $PI^-$ cells | |
|---|---|---|
| **Mice** | **$CD31^-CD45^-Ter119^-$** | **$CD31^+CD45^+Ter119^+$** |
| *Ngf*$^{fl/fl}$ | 35.4 | 64.6 |
| *Ngf*$^{LysM}$ | 31.0 | 69.0 |

**Table S2.** Differentially expressed genes of cluster 5 from $p75^{PDGFR\alpha}$ mice in comparison to $p75^{fl/fl}$ mice.

| Gene | avg_logFC | p_val_adj | Cluster |
|---|---|---|---|
| Gm26917 | 2.756371 | 4.23E-79 | 5 |
| Malat1 | 2.04657844 | 6.29E-79 | 5 |
| Lars2 | 1.75771496 | 1.29E-67 | 5 |
| Meg3 | 1.65886322 | 1.19E-30 | 5 |
| Xist | 1.60505268 | 2.57E-33 | 5 |
| Gm42418 | 1.46692346 | 2.32E-53 | 5 |
| mt-Atp6 | 1.46556536 | 1.83E-88 | 5 |
| mt-Co2 | 1.4210027 | 5.11E-91 | 5 |
| mt-Co1 | 1.41561465 | 1.04E-87 | 5 |
| mt-Co3 | 1.41325514 | 8.68E-89 | 5 |
| Rian | 1.40055979 | 2.34E-30 | 5 |
| mt-Nd1 | 1.33077477 | 9.88E-83 | 5 |
| mt-Nd5 | 1.27996765 | 7.26E-59 | 5 |
| mt-Cytb | 1.27131161 | 9.22E-82 | 5 |
| mt-Nd4 | 1.26767758 | 2.15E-81 | 5 |
| mt-Nd3 | 1.2539379 | 3.10E-65 | 5 |
| mt-Nd2 | 1.237207 | 2.14E-73 | 5 |
| Kcnq1ot1 | 1.22357867 | 4.22E-24 | 5 |
| Neat1 | 1.16267805 | 7.80E-34 | 5 |
| Dnm3os | 1.0914986 | 1.08E-35 | 5 |
| mt-Nd4l | 0.95935865 | 2.87E-41 | 5 |
| Nktr | 0.95208603 | 2.86E-44 | 5 |
| AY036118 | 0.93058927 | 1.08E-44 | 5 |
| mt-Atp8 | 0.92246563 | 1.53E-38 | 5 |
| Col7a1 | 0.91205676 | 9.26E-18 | 5 |
| Dst | 0.89194034 | 8.61E-49 | 5 |
| Col5a3 | 0.87470159 | 3.19E-12 | 5 |
| B830012L14Rik | 0.87459019 | 4.32E-25 | 5 |
| Col5a1 | 0.85947061 | 1.19E-47 | 5 |
| Ogt | 0.8432767 | 1.97E-31 | 5 |
| Col27a1 | 0.79872506 | 1.41E-30 | 5 |
| Ccnl2 | 0.79807569 | 9.46E-31 | 5 |
| AC149090.1 | 0.79620379 | 5.27E-21 | 5 |
| Krit1 | 0.79607439 | 2.25E-28 | 5 |
| Mast4 | 0.7956282 | 4.30E-31 | 5 |
| Mir99ahg | 0.79091837 | 3.76E-19 | 5 |
| Dleu2 | 0.78935654 | 3.66E-15 | 5 |

| | | | |
|---|---|---|---|
| Hjurp | 0.77515591 | 1.94E-25 | 5 |
| Col6a3 | 0.76887731 | 1.70E-24 | 5 |
| Ddx17 | 0.75813096 | 6.89E-29 | 5 |
| Nfat5 | 0.75662507 | 3.66E-25 | 5 |
| Ebf1 | 0.74821591 | 2.89E-08 | 5 |
| Pnisr | 0.74436579 | 1.44E-33 | 5 |
| Gm47283 | 0.73182908 | 4.13E-22 | 5 |
| Hspg2 | 0.73114439 | 1.09E-33 | 5 |
| Tnc | 0.72909432 | 2.68E-06 | 5 |
| Col16a1 | 0.72653922 | 1.60E-26 | 5 |
| Luc7l2 | 0.72344066 | 1.68E-28 | 5 |
| Zc3h7a | 0.72163691 | 5.81E-25 | 5 |
| Col12a1 | 0.71787243 | 3.01E-10 | 5 |
| Map4k4 | 0.71188242 | 2.08E-30 | 5 |
| Ttc14 | 0.71043316 | 3.31E-22 | 5 |
| Son | 0.69371176 | 7.09E-34 | 5 |
| Nfkb1 | 0.67480906 | 7.78E-10 | 5 |
| Adam33 | 0.67439662 | 1.52E-11 | 5 |
| Fus | 0.67032939 | 1.32E-26 | 5 |
| Uhrf2 | 0.6675981 | 2.71E-19 | 5 |
| Ankrd11 | 0.66489719 | 1.20E-34 | 5 |
| Hivep2 | 0.65925291 | 6.71E-19 | 5 |
| Snrnp70 | 0.65771331 | 2.70E-38 | 5 |
| Trio | 0.65284247 | 6.10E-12 | 5 |
| Hmcn1 | 0.64818791 | 2.22E-16 | 5 |
| Ubn2 | 0.64541688 | 4.14E-23 | 5 |
| Hnrnph1 | 0.641747 | 3.13E-30 | 5 |
| Ccnl1 | 0.62421505 | 1.60E-14 | 5 |
| Fbn2 | 0.62131287 | 2.15E-16 | 5 |
| Syne1 | 0.62119331 | 7.07E-14 | 5 |
| Col1a1 | 0.62007939 | 1.13E-34 | 5 |
| Nisch | 0.6151973 | 1.55E-34 | 5 |
| Dennd4a | 0.61506993 | 7.59E-06 | 5 |
| Arglu1 | 0.60775911 | 4.68E-24 | 5 |
| Zcchc7 | 0.60569117 | 5.47E-21 | 5 |
| Fbln2 | 0.6044937 | 5.35E-19 | 5 |
| Lrp1 | 0.60417832 | 2.69E-24 | 5 |
| Rsrp1 | 0.60320165 | 7.38E-32 | 5 |
| Mrc2 | 0.58740299 | 3.37E-20 | 5 |
| Zfc3h1 | 0.58660363 | 1.26E-25 | 5 |
| Mirg | 0.58572597 | 8.98E-09 | 5 |
| Kdm6b | 0.58363154 | 9.73E-06 | 5 |

| | | | |
|---|---|---|---|
| Ankrd17 | 0.58194012 | 1.86E-19 | 5 |
| Mir100hg | 0.57522676 | 3.89E-11 | 5 |
| Lamb1 | 0.5735042 | 1.19E-18 | 5 |
| Prpf4b | 0.57343216 | 1.50E-11 | 5 |
| Sbno2 | 0.57246861 | 7.04E-12 | 5 |
| Thbs2 | 0.57020493 | 7.67E-13 | 5 |
| Dot1l | 0.56640151 | 7.57E-08 | 5 |
| Mycbp2 | 0.55979005 | 1.22E-19 | 5 |
| Ankrd12 | 0.55972026 | 3.56E-11 | 5 |
| Stk38 | 0.55873819 | 2.55E-11 | 5 |
| Plec | 0.55675213 | 1.33E-14 | 5 |
| Pkd2 | 0.55545763 | 1.46E-16 | 5 |
| Srek1 | 0.55267944 | 3.42E-23 | 5 |
| Taf1d | 0.55263512 | 1.96E-12 | 5 |
| Akap13 | 0.55243711 | 2.03E-23 | 5 |
| Runx1 | 0.55077252 | 6.05E-14 | 5 |
| Golga4 | 0.54764063 | 6.56E-17 | 5 |
| Clk1 | 0.54743611 | 8.15E-21 | 5 |
| Emilin1 | 0.54689066 | 5.39E-14 | 5 |
| Kmt2a | 0.54550861 | 3.32E-20 | 5 |
| Hhipl1 | 0.54351442 | 2.20E-13 | 5 |
| Srsf11 | 0.54206337 | 3.25E-26 | 5 |
| Vmp1 | 0.53776068 | 1.16E-14 | 5 |
| Tia1 | 0.53727207 | 2.03E-05 | 5 |
| Clec2d | 0.5370208 | 0.0134371 | 5 |
| Tnrc6c | 0.53606264 | 1.15E-17 | 5 |
| Zbtb20 | 0.53575989 | 6.92E-11 | 5 |
| Col5a2 | 0.5351243 | 5.57E-27 | 5 |
| Utrn | 0.52793892 | 7.49E-17 | 5 |
| Creb5 | 0.52613052 | 0.04452381 | 5 |
| Aebp1 | 0.5257943 | 1.05E-11 | 5 |
| Srsf5 | 0.52565743 | 7.22E-25 | 5 |
| Mmp14 | 0.52335152 | 5.00E-16 | 5 |
| Spag9 | 0.52160722 | 4.43E-23 | 5 |
| Fgfr1 | 0.51970256 | 1.08E-15 | 5 |
| Fubp1 | 0.51811868 | 9.27E-16 | 5 |
| Phip | 0.51652202 | 5.80E-12 | 5 |
| Prpf38b | 0.51556844 | 2.28E-21 | 5 |
| Bclaf1 | 0.51398439 | 2.98E-22 | 5 |
| Hk2 | 0.51284051 | 0.00026599 | 5 |
| Zfp292 | 0.51093197 | 2.04E-12 | 5 |
| Brwd1 | 0.50881 | 1.68E-20 | 5 |

| | | | |
|---|---|---|---|
| Zeb1 | 0.50872949 | 6.07E-22 | 5 |
| Auts2 | 0.50780947 | 1.58E-13 | 5 |
| Slc38a10 | 0.50773229 | 1.19E-15 | 5 |
| Prpf39 | 0.50144368 | 3.51E-10 | 5 |
| Thbs3 | 0.49940571 | 9.45E-17 | 5 |
| Mafg | 0.49829316 | 6.64E-18 | 5 |
| Gpc1 | 0.49317568 | 1.39E-07 | 5 |
| Setd5 | 0.49000788 | 5.28E-14 | 5 |
| Etnk1 | 0.48712682 | 5.70E-08 | 5 |
| Golgb1 | 0.48632862 | 9.58E-17 | 5 |
| Qsox1 | 0.48604324 | 1.61E-17 | 5 |
| Ptprs | 0.48094558 | 1.55E-18 | 5 |
| Akap9 | 0.47940837 | 2.07E-13 | 5 |
| Mdm4 | 0.47800825 | 1.81E-07 | 5 |
| Pxdn | 0.47778267 | 5.86E-11 | 5 |
| mt-Nd6 | 0.47498325 | 2.82E-09 | 5 |
| Ahnak2 | 0.47483826 | 0.00010124 | 5 |
| Sox4 | 0.47434035 | 1.17E-06 | 5 |
| Slc20a1 | 0.47296797 | 7.88E-06 | 5 |
| Cspp1 | 0.47152172 | 1.03E-08 | 5 |
| Zeb2 | 0.46989491 | 4.11E-16 | 5 |
| Ccnt2 | 0.46926008 | 2.57E-09 | 5 |
| Ash1l | 0.46844964 | 2.28E-17 | 5 |
| Nsd3 | 0.46735843 | 6.69E-14 | 5 |
| Chd2 | 0.46711647 | 2.10E-07 | 5 |
| Rbm39 | 0.46621757 | 3.88E-17 | 5 |
| Zfhx4 | 0.4662054 | 1.33E-08 | 5 |
| Adamts14 | 0.46586748 | 2.04E-07 | 5 |
| Gls | 0.45929657 | 9.15E-13 | 5 |
| Psd3 | 0.45795506 | 5.42E-06 | 5 |
| Fn1 | 0.4572526 | 1.95E-11 | 5 |
| Alkbh1 | 0.45649894 | 4.33E-10 | 5 |
| Camk2d | 0.45571853 | 3.88E-11 | 5 |
| Pkd1 | 0.45335371 | 6.84E-15 | 5 |
| Iqgap1 | 0.45252501 | 1.70E-12 | 5 |
| Slc39a14 | 0.45233148 | 8.65E-07 | 5 |
| Mkln1 | 0.45224828 | 6.20E-10 | 5 |
| Ptprd | 0.45223803 | 3.25E-06 | 5 |
| Bmp1 | 0.45160926 | 4.99E-14 | 5 |
| Rrbp1 | 0.45031385 | 4.21E-14 | 5 |
| Crebzf | 0.45020593 | 0.00691361 | 5 |
| Miat | 0.44973242 | 2.35E-07 | 5 |

| | | | |
|---|---|---|---|
| Mapk8ip3 | 0.44966567 | 3.70E-07 | 5 |
| Safb2 | 0.44853797 | 1.12E-10 | 5 |
| Bnc2 | 0.44826367 | 1.10E-06 | 5 |
| Atp2b4 | 0.44824173 | 0.00086053 | 5 |
| Tet2 | 0.44792074 | 3.49E-06 | 5 |
| Tra2a | 0.44727928 | 6.55E-10 | 5 |
| Aebp2 | 0.44614265 | 3.19E-13 | 5 |
| Acin1 | 0.44532558 | 9.60E-17 | 5 |
| Arhgap31 | 0.44391387 | 5.14E-05 | 5 |
| Vcan | 0.44335481 | 1.76E-07 | 5 |
| Tcf4 | 0.44329151 | 1.62E-09 | 5 |
| Gm20559 | 0.44253838 | 2.80E-06 | 5 |
| Klhl24 | 0.44210242 | 2.57E-06 | 5 |
| Maf | 0.44147851 | 3.28E-06 | 5 |
| Mbnl2 | 0.43933639 | 2.58E-14 | 5 |
| Nbeal1 | 0.43903051 | 2.47E-10 | 5 |
| Nfatc4 | 0.43882273 | 4.01E-10 | 5 |
| Zfhx3 | 0.43846831 | 4.85E-13 | 5 |
| Abcc5 | 0.43687548 | 1.02E-06 | 5 |
| Cuedc1 | 0.43685798 | 1.02E-12 | 5 |
| Setd2 | 0.43658846 | 4.22E-06 | 5 |
| Mical2 | 0.43495204 | 3.70E-07 | 5 |
| Rbm5 | 0.43476185 | 6.68E-15 | 5 |
| Sorcs2 | 0.4339988 | 1.21E-07 | 5 |
| Mga | 0.43339967 | 4.91E-09 | 5 |
| Col1a2 | 0.43242673 | 9.89E-14 | 5 |
| Phf3 | 0.43131658 | 2.20E-16 | 5 |
| Ncam1 | 0.43108607 | 9.35E-10 | 5 |
| Pdgfrb | 0.43028378 | 5.83E-05 | 5 |
| Dlg4 | 0.4288977 | 5.20E-08 | 5 |
| Rc3h2 | 0.42802966 | 8.03E-11 | 5 |
| Tcirg1 | 0.42739041 | 2.10E-06 | 5 |
| Herc1 | 0.42497848 | 1.71E-07 | 5 |
| Rictor | 0.42455221 | 7.62E-07 | 5 |
| Gtpbp2 | 0.42432634 | 4.63E-05 | 5 |
| Rock2 | 0.42243414 | 2.05E-13 | 5 |
| Atrx | 0.4214536 | 9.17E-11 | 5 |
| Zfp451 | 0.42076888 | 2.53E-08 | 5 |
| Gfpt1 | 0.42063753 | 3.92E-09 | 5 |
| Nrp2 | 0.42041437 | 2.10E-08 | 5 |
| Macf1 | 0.41831939 | 1.46E-10 | 5 |
| Dtx3 | 0.41691854 | 7.18E-06 | 5 |

| | | | |
|---|---|---|---|
| Tead1 | 0.41637937 | 1.66E-08 | 5 |
| Rai1 | 0.41533245 | 1.08E-05 | 5 |
| Resf1 | 0.41464513 | 1.08E-05 | 5 |
| Nav3 | 0.41455721 | 1.37E-08 | 5 |
| Cdk14 | 0.41431637 | 1.47E-08 | 5 |
| Edil3 | 0.4139536 | 5.10E-06 | 5 |
| Furin | 0.41350368 | 5.59E-09 | 5 |
| Slc38a6 | 0.41319777 | 1.92E-12 | 5 |
| Cdk12 | 0.4123516 | 7.15E-06 | 5 |
| Kmt2e | 0.41164729 | 1.14E-15 | 5 |
| Safb | 0.40993983 | 2.02E-08 | 5 |
| Bptf | 0.40966941 | 6.34E-13 | 5 |
| Piezo1 | 0.40855896 | 9.47E-07 | 5 |
| Igsf3 | 0.40716872 | 0.00123155 | 5 |
| Glg1 | 0.4063901 | 8.28E-13 | 5 |
| Tanc1 | 0.40499331 | 6.95E-05 | 5 |
| Pdgfra | 0.40470747 | 0.00032691 | 5 |
| Stx16 | 0.4038559 | 0.00019419 | 5 |
| Atp2b1 | 0.4036764 | 3.26E-10 | 5 |
| Baiap2 | 0.40290382 | 0.02537364 | 5 |
| Lrrc15 | 0.40228188 | 1.30E-05 | 5 |
| Plod2 | 0.40174747 | 2.26E-09 | 5 |
| Scarf2 | 0.39827102 | 3.31E-05 | 5 |
| Nemf | 0.39758747 | 1.50E-09 | 5 |
| Jmjd1c | 0.39729749 | 1.29E-07 | 5 |
| Smc5 | 0.39726925 | 1.14E-06 | 5 |
| Atxn1 | 0.39561394 | 6.28E-08 | 5 |
| Ddx50 | 0.39527075 | 3.40E-11 | 5 |
| Prrc2c | 0.394378 | 3.47E-15 | 5 |
| Srrm2 | 0.39367216 | 4.19E-15 | 5 |
| Ttc3 | 0.39158536 | 1.88E-17 | 5 |
| Dlc1 | 0.39155728 | 7.84E-08 | 5 |
| Clk4 | 0.39073603 | 4.28E-07 | 5 |
| Zcchc14 | 0.39067903 | 0.00663432 | 5 |
| Enah | 0.39028677 | 4.59E-09 | 5 |
| Lrig3 | 0.38615387 | 2.79E-06 | 5 |
| Chd3 | 0.38539292 | 1.54E-06 | 5 |
| Zmym5 | 0.38511422 | 0.00100633 | 5 |
| Nav1 | 0.38501928 | 1.04E-05 | 5 |
| Gigyf1 | 0.38451979 | 2.80E-05 | 5 |
| Raph1 | 0.38336311 | 0.01340591 | 5 |
| Myh9 | 0.3832309 | 1.45E-08 | 5 |

| | | | |
|---|---|---|---|
| Sh3pxd2a | 0.38298348 | 1.53E-09 | 5 |
| Ago3 | 0.38139288 | 9.35E-07 | 5 |
| Rsrc2 | 0.38135048 | 1.22E-11 | 5 |
| Adamts10 | 0.38078567 | 2.92E-06 | 5 |
| Abcd4 | 0.3791896 | 1.79E-07 | 5 |
| Kdm5a | 0.37779891 | 0.00017129 | 5 |
| Tln2 | 0.37760487 | 5.55E-09 | 5 |
| Cdc42bpa | 0.377415 | 2.35E-08 | 5 |
| Zfp950 | 0.37710946 | 0.00077953 | 5 |
| Parp8 | 0.37584814 | 1.95E-06 | 5 |
| Igf1r | 0.37522767 | 0.00392169 | 5 |
| Tacc1 | 0.37468551 | 2.52E-05 | 5 |
| Usp34 | 0.37450809 | 2.02E-09 | 5 |
| Rbms3 | 0.37438457 | 3.53E-05 | 5 |
| Arhgef40 | 0.37433297 | 3.17E-06 | 5 |
| Gpatch8 | 0.37407665 | 0.00043535 | 5 |
| Tnrc6b | 0.37281253 | 1.45E-07 | 5 |
| Gatad2b | 0.37088555 | 3.29E-08 | 5 |
| 4632427E13Rik | 0.3707821 | 0.00612754 | 5 |
| Tasor | 0.36944232 | 0.03781291 | 5 |
| Fam193b | 0.36937768 | 3.50E-09 | 5 |
| Khdc4 | 0.36886896 | 5.03E-05 | 5 |
| Apaf1 | 0.36867633 | 0.00165493 | 5 |
| Plekhg2 | 0.36864121 | 0.00012602 | 5 |
| Ptgfrn | 0.3672523 | 2.27E-08 | 5 |
| Smg1 | 0.36655226 | 1.33E-08 | 5 |
| Cpsf6 | 0.36576587 | 1.77E-08 | 5 |
| Tsix | 0.36538421 | 1.63E-21 | 5 |
| Sfpq | 0.36423466 | 1.43E-13 | 5 |
| Tnrc6a | 0.36380251 | 2.33E-07 | 5 |
| Phldb1 | 0.36341689 | 0.00041107 | 5 |
| Col3a1 | 0.36312729 | 5.53E-09 | 5 |
| Zfp644 | 0.36221377 | 1.61E-08 | 5 |
| Rb1cc1 | 0.36220089 | 1.69E-09 | 5 |
| Sbno1 | 0.36186273 | 1.29E-10 | 5 |
| Tmcc1 | 0.36131524 | 0.00013896 | 5 |
| Sf3b3 | 0.36026807 | 1.46E-07 | 5 |
| Paxbp1 | 0.3599685 | 0.00014645 | 5 |
| Airn | 0.35990801 | 2.04E-06 | 5 |
| Nsd1 | 0.35774821 | 8.58E-10 | 5 |
| Igf2r | 0.35734545 | 2.13E-10 | 5 |
| Pvr | 0.3568167 | 0.00096055 | 5 |

| | | | |
|---|---|---|---|
| Tmem259 | 0.35617179 | 8.28E-05 | 5 |
| Ranbp2 | 0.35587207 | 5.79E-08 | 5 |
| Htt | 0.35587151 | 2.40E-05 | 5 |
| Arhgap17 | 0.35542967 | 0.00587021 | 5 |
| Ano6 | 0.35518454 | 0.00153934 | 5 |
| Dync1h1 | 0.35484772 | 2.06E-14 | 5 |
| Fnip1 | 0.35144751 | 3.16E-06 | 5 |
| Fxr2 | 0.35127907 | 0.00033254 | 5 |
| P3h1 | 0.35060998 | 2.73E-07 | 5 |
| Csnk1g1 | 0.35035104 | 0.00011321 | 5 |
| Med13l | 0.34996438 | 4.22E-07 | 5 |
| Cdip1 | 0.34992267 | 7.92E-05 | 5 |
| Ascc3 | 0.34686076 | 2.40E-08 | 5 |
| Dnmt3a | 0.34643354 | 9.77E-06 | 5 |
| R3hdm1 | 0.34631176 | 1.41E-06 | 5 |
| Adgrl1 | 0.34620875 | 3.91E-07 | 5 |
| Plekha1 | 0.34584923 | 0.00123658 | 5 |
| Atp13a3 | 0.34576613 | 0.00032996 | 5 |
| Mbtd1 | 0.34551397 | 0.01940742 | 5 |
| Far1 | 0.34398102 | 1.89E-05 | 5 |
| Flnc | 0.34362403 | 0.0004518 | 5 |
| Tial1 | 0.34320967 | 8.96E-07 | 5 |
| Ncln | 0.3422295 | 0.00045202 | 5 |
| Tns1 | 0.34158793 | 3.73E-05 | 5 |
| Parp3 | 0.34122338 | 0.00023017 | 5 |
| 5031425E22Rik | 0.33830541 | 0.01440726 | 5 |
| Esyt2 | 0.3381333 | 5.03E-08 | 5 |
| Leng8 | 0.33771286 | 6.73E-06 | 5 |
| D430042O09Rik | 0.3370581 | 0.01698492 | 5 |
| Riok3 | 0.33661737 | 1.16E-06 | 5 |
| Birc6 | 0.33635143 | 5.54E-13 | 5 |
| Trip12 | 0.33562586 | 9.07E-08 | 5 |
| Gpr153 | 0.33530976 | 1.24E-07 | 5 |
| Tcf12 | 0.33488404 | 4.80E-08 | 5 |
| Atxn2 | 0.33412791 | 4.65E-05 | 5 |
| Ktn1 | 0.334006 | 6.54E-09 | 5 |
| Oga | 0.33387583 | 8.63E-05 | 5 |
| Fbn1 | 0.33381967 | 3.04E-05 | 5 |
| Nxf1 | 0.33333494 | 0.0040879 | 5 |
| Adamts2 | 0.33328423 | 1.78E-06 | 5 |
| Sf3b1 | 0.33314678 | 2.42E-14 | 5 |
| Npepps | 0.33265456 | 1.38E-05 | 5 |

| | | | |
|---|---|---|---|
| Brd8 | 0.33193989 | 0.0023908 | 5 |
| Wsb1 | 0.33171251 | 7.77E-06 | 5 |
| Adam12 | 0.33167717 | 9.71E-09 | 5 |
| Adamts4 | 0.33073826 | 0.0029114 | 5 |
| Skil | 0.33067085 | 0.01126941 | 5 |
| Copa | 0.33047203 | 1.67E-13 | 5 |
| Adamtsl1 | 0.33043919 | 2.59E-06 | 5 |
| Plxnb2 | 0.3303261 | 2.81E-07 | 5 |
| Kdm2a | 0.32997164 | 7.24E-09 | 5 |
| Ankrd16 | 0.32956364 | 0.00010063 | 5 |
| Loxl3 | 0.32875414 | 6.13E-06 | 5 |
| Osmr | 0.32848522 | 0.00026047 | 5 |
| Cd2ap | 0.32748595 | 1.75E-07 | 5 |
| Lzts2 | 0.32726246 | 0.00025905 | 5 |
| Mib1 | 0.32694968 | 4.16E-08 | 5 |
| Map3k2 | 0.32686612 | 0.00092792 | 5 |
| Kif26b | 0.32673049 | 0.00483963 | 5 |
| Cyth3 | 0.32581946 | 0.02053363 | 5 |
| Rbfox2 | 0.32564391 | 3.94E-10 | 5 |
| Chd9 | 0.32525512 | 2.43E-05 | 5 |
| Myo9b | 0.32514694 | 4.00E-05 | 5 |
| Zfp516 | 0.32460344 | 5.94E-06 | 5 |
| Supt20 | 0.32389898 | 0.00038584 | 5 |
| Tulp4 | 0.32366386 | 3.51E-06 | 5 |
| Pds5a | 0.32349799 | 0.00062176 | 5 |
| Sptbn1 | 0.32330225 | 3.29E-09 | 5 |
| Map4 | 0.32304867 | 1.10E-12 | 5 |
| 4932438A13Rik | 0.32296552 | 0.00384767 | 5 |
| Adgra2 | 0.32275109 | 0.00030094 | 5 |
| Fnbp4 | 0.32216434 | 0.0255764 | 5 |
| Zfp266 | 0.3214742 | 0.00071367 | 5 |
| Rbm25 | 0.32131538 | 2.14E-09 | 5 |
| Adcy7 | 0.32126553 | 0.00078432 | 5 |
| Mapk6 | 0.32053913 | 1.72E-05 | 5 |
| Itch | 0.32038638 | 0.01723053 | 5 |
| Ppp1r12a | 0.3203692 | 0.00014024 | 5 |
| Ecpas | 0.31986205 | 3.66E-06 | 5 |
| Tardbp | 0.31891284 | 6.69E-10 | 5 |
| Trrap | 0.31880731 | 3.19E-06 | 5 |
| Fto | 0.31849533 | 0.0002826 | 5 |
| Tsc1 | 0.31838744 | 0.00145749 | 5 |
| Ewsr1 | 0.31769642 | 2.75E-10 | 5 |

| | | | |
|---|---|---|---|
| Kif1b | 0.31705166 | 7.75E-08 | 5 |
| Mon2 | 0.31689608 | 3.54E-05 | 5 |
| Celf1 | 0.31624459 | 5.13E-08 | 5 |
| Taok1 | 0.31566755 | 3.48E-08 | 5 |
| Zfp871 | 0.31545712 | 0.03637204 | 5 |
| Arhgef25 | 0.31537288 | 0.00635054 | 5 |
| Dvl1 | 0.31412498 | 0.03792169 | 5 |
| Mark3 | 0.31358006 | 0.01677052 | 5 |
| Trim35 | 0.31344272 | 0.00081744 | 5 |
| Selenoi | 0.3127468 | 0.001111 | 5 |
| Lamc1 | 0.31267484 | 6.61E-07 | 5 |
| Flna | 0.31241607 | 7.97E-06 | 5 |
| Ccdc80 | 0.31235069 | 0.00050499 | 5 |
| Tra2b | 0.31228789 | 5.65E-08 | 5 |
| Huwe1 | 0.31208098 | 4.80E-07 | 5 |
| Chd4 | 0.31189069 | 0.00498769 | 5 |
| Lama4 | 0.31171529 | 6.28E-07 | 5 |
| Asph | 0.31114549 | 1.06E-06 | 5 |
| Tanc2 | 0.31107187 | 0.00214024 | 5 |
| Fosl2 | 0.31105393 | 0.00247793 | 5 |
| Sec61a1 | 0.3109141 | 5.84E-06 | 5 |
| Emsy | 0.31082144 | 0.00051061 | 5 |
| Cnot1 | 0.31041314 | 0.00103552 | 5 |
| Usp33 | 0.31040749 | 0.00384757 | 5 |
| Srcap | 0.31007469 | 0.02079977 | 5 |
| Usp36 | 0.30976265 | 0.00101624 | 5 |
| Itga5 | 0.30950205 | 0.00049298 | 5 |
| Zzef1 | 0.30914016 | 1.60E-06 | 5 |
| Sec24a | 0.30907382 | 0.00060552 | 5 |
| Esyt1 | 0.30884028 | 1.20E-07 | 5 |
| Luc7l3 | 0.30875113 | 0.02121635 | 5 |
| Trip11 | 0.30859225 | 0.00010476 | 5 |
| Aff4 | 0.30848495 | 8.94E-07 | 5 |
| Atxn2l | 0.30848426 | 0.00040776 | 5 |
| Tnrc18 | 0.3083422 | 0.00010637 | 5 |
| Ppfibp1 | 0.30718136 | 3.35E-09 | 5 |
| Tsc22d2 | 0.30706079 | 0.00093104 | 5 |
| Nufip2 | 0.30677909 | 6.91E-05 | 5 |
| Otud7b | 0.30659616 | 0.00184077 | 5 |
| Loxl2 | 0.30642371 | 0.00815747 | 5 |
| Cep350 | 0.30633369 | 0.00618916 | 5 |
| Zfp9 | 0.30580209 | 0.01824914 | 5 |

| | | | |
|---|---|---|---|
| Igsf10 | 0.30524303 | 0.0001468 | 5 |
| Tgfb2 | 0.30512408 | 0.00034313 | 5 |
| Qk | 0.30507502 | 1.35E-07 | 5 |
| Snrnp48 | 0.3044494 | 9.55E-05 | 5 |
| Hook3 | 0.30329578 | 0.00054446 | 5 |
| Rlf | 0.30290975 | 0.00528614 | 5 |
| Rbbp6 | 0.30264653 | 1.56E-07 | 5 |
| Kmt2c | 0.30094041 | 0.00187328 | 5 |
| Csnk1a1 | 0.30032716 | 0.00012165 | 5 |
| Prrc1 | 0.30004486 | 4.36E-05 | 5 |
| Strn3 | 0.29977962 | 3.02E-06 | 5 |
| Chpf2 | 0.29939876 | 0.00130563 | 5 |
| Olfml2b | 0.29892614 | 1.66E-05 | 5 |
| Gnl3 | 0.29846318 | 2.91E-07 | 5 |
| P3h3 | 0.29816267 | 7.63E-08 | 5 |
| Ubr2 | 0.29731454 | 8.40E-05 | 5 |
| Ikbkb | 0.29729166 | 0.02283246 | 5 |
| Hnrnpdl | 0.29692101 | 1.45E-08 | 5 |
| Wwc2 | 0.29691751 | 1.25E-06 | 5 |
| Tcf7l2 | 0.29503507 | 0.04728663 | 5 |
| Tut7 | 0.2949352 | 0.00187274 | 5 |
| Flnb | 0.29473291 | 0.00107347 | 5 |
| Lpp | 0.29470106 | 8.48E-05 | 5 |
| Zfp469 | 0.2946528 | 6.09E-05 | 5 |
| Inf2 | 0.29400177 | 0.00088764 | 5 |
| Aff1 | 0.29389066 | 0.00360293 | 5 |
| Atp2a2 | 0.2929597 | 2.64E-09 | 5 |
| Pip5k1a | 0.29189944 | 0.00018672 | 5 |
| Phf21a | 0.29177657 | 0.04297569 | 5 |
| Ythdc1 | 0.29059523 | 0.0108109 | 5 |
| Aak1 | 0.28974145 | 4.33E-07 | 5 |
| Sema6c | 0.28783943 | 0.00412446 | 5 |
| Ephb2 | 0.28623473 | 0.02789639 | 5 |
| Tmx3 | 0.2847686 | 0.01103858 | 5 |
| Hdac7 | 0.28448791 | 0.00660269 | 5 |
| Antxr1 | 0.28400008 | 0.00146086 | 5 |
| Srrt | 0.2834098 | 0.00215584 | 5 |
| Stx5a | 0.28309913 | 0.00651834 | 5 |
| Ubr5 | 0.28304889 | 6.63E-06 | 5 |
| Arhgef1 | 0.28229093 | 0.01641294 | 5 |
| Lnpep | 0.28147292 | 3.71E-05 | 5 |
| Tnks2 | 0.28146287 | 0.00184931 | 5 |

| | | | |
|---|---|---|---|
| Zfp280d | 0.2811803 | 0.02640616 | 5 |
| Prrc2a | 0.28028622 | 1.86E-06 | 5 |
| Zkscan3 | 0.27921746 | 0.02599017 | 5 |
| Pcdh19 | 0.27819031 | 0.0260666 | 5 |
| Pkn2 | 0.27815813 | 0.00096326 | 5 |
| Kidins220 | 0.278117 | 0.04344644 | 5 |
| Sulf1 | 0.27735357 | 0.00066509 | 5 |
| Dmtf1 | 0.27633887 | 0.00299254 | 5 |
| Yeats2 | 0.27606484 | 0.00041002 | 5 |
| Col6a1 | 0.27596353 | 0.00971932 | 5 |
| Sh3pxd2b | 0.27592706 | 9.11E-06 | 5 |
| Nipbl | 0.27395603 | 1.03E-05 | 5 |
| Hnrnpl | 0.27353729 | 0.00014702 | 5 |
| Megf8 | 0.27346012 | 0.00090644 | 5 |
| Gapvd1 | 0.27245828 | 0.00734341 | 5 |
| Itgav | 0.27187141 | 0.00040734 | 5 |
| Fam13b | 0.27168745 | 6.34E-05 | 5 |
| Pura | 0.26778044 | 1.36E-09 | 5 |
| Ahnak | 0.26708278 | 0.00058204 | 5 |
| Frmd6 | 0.26690628 | 5.07E-06 | 5 |
| Cul7 | 0.26584063 | 0.00068515 | 5 |
| Baz1b | 0.26441283 | 0.00718548 | 5 |
| Lncpint | 0.26400623 | 0.01353226 | 5 |
| Ncor1 | 0.26305571 | 2.30E-07 | 5 |
| Brd1 | 0.2614468 | 1.80E-05 | 5 |
| Fkbp9 | 0.26101817 | 0.00142806 | 5 |
| Ccdc88a | 0.26098466 | 2.82E-05 | 5 |
| Zswim8 | 0.26092073 | 0.03042535 | 5 |
| Fndc3b | 0.25993733 | 4.85E-07 | 5 |
| Dnm1 | 0.25659593 | 0.02117681 | 5 |
| Xrn1 | 0.25446595 | 0.00321451 | 5 |
| Tm9sf4 | 0.25228585 | 0.00027283 | 5 |
| Gcn1 | 0.25182971 | 0.00102993 | 5 |
| BC005537 | 0.25056039 | 1.40E-05 | 5 |
| Dhx9 | 0.25031247 | 0.01527382 | 5 |

**Table S3.** Differentially expressed genes of cluster 6 from $p75^{PDGFR\alpha}$ mice in comparison to $p75^{fl/fl}$ mice.

| Gene | avg_logFC | p_val_adj | Cluster |
|---|---|---|---|
| S100a9 | 2.78970369 | 1.09E-65 | 6 |
| S100a8 | 2.7457214 | 2.80E-74 | 6 |
| Ngp | 2.29442596 | 1.12E-46 | 6 |
| Camp | 2.25425912 | 1.48E-23 | 6 |
| Lyz2 | 2.24038072 | 3.08E-168 | 6 |
| Retnlg | 2.01821224 | 1.51E-56 | 6 |
| Lcn2 | 1.95546791 | 7.78E-67 | 6 |
| Tyrobp | 1.68114859 | 2.49E-176 | 6 |
| Fcer1g | 1.62003506 | 3.19E-183 | 6 |
| Pf4 | 1.51338472 | 3.39E-38 | 6 |
| Cd74 | 1.51286045 | 3.36E-21 | 6 |
| Ctss | 1.36982733 | 9.75E-66 | 6 |
| Ltf | 1.36326356 | 1.98E-39 | 6 |
| Cd52 | 1.33075535 | 1.24E-186 | 6 |
| Apoe | 1.29982021 | 1.67E-35 | 6 |
| Wfdc21 | 1.29841443 | 4.46E-87 | 6 |
| Il1b | 1.23016989 | 1.20E-54 | 6 |
| C1qa | 1.1972231 | 3.50E-32 | 6 |
| Cxcl2 | 1.07291696 | 9.30E-41 | 6 |
| C1qb | 1.06762843 | 6.02E-37 | 6 |
| Srgn | 1.06670758 | 1.22E-76 | 6 |
| Slpi | 1.05777133 | 4.32E-79 | 6 |
| Alox5ap | 1.05004338 | 1.72E-169 | 6 |
| Cd14 | 1.04522715 | 5.69E-61 | 6 |
| Lgals3 | 1.04292347 | 1.27E-53 | 6 |
| Laptm5 | 1.02023456 | 1.11E-162 | 6 |
| H2-Aa | 1.0124058 | 6.54E-10 | 6 |
| Pglyrp1 | 1.00361398 | 2.34E-133 | 6 |
| Coro1a | 0.99085463 | 1.70E-169 | 6 |
| C1qc | 0.9456425 | 3.39E-51 | 6 |
| Mmp8 | 0.93401328 | 1.05E-118 | 6 |
| Lcp1 | 0.93009676 | 2.06E-202 | 6 |
| Ifitm6 | 0.91715532 | 2.95E-82 | 6 |
| H2-Ab1 | 0.87707585 | 6.10E-05 | 6 |
| Ccl6 | 0.86011353 | 3.71E-105 | 6 |
| Chil3 | 0.82820684 | 1.66E-42 | 6 |
| Cybb | 0.82497413 | 4.73E-200 | 6 |

| | | | |
|---|---|---|---|
| Hp | 0.80626693 | 8.13E-79 | 6 |
| Cd68 | 0.7898913 | 7.38E-73 | 6 |
| Cyba | 0.78673482 | 1.53E-47 | 6 |
| Ucp2 | 0.77216351 | 3.84E-93 | 6 |
| Rac2 | 0.7659813 | 2.58E-201 | 6 |
| Ccl9 | 0.72738926 | 1.82E-54 | 6 |
| Psap | 0.72164727 | 5.62E-31 | 6 |
| Slfn2 | 0.71463745 | 2.46E-110 | 6 |
| Mmp9 | 0.7135716 | 1.13E-36 | 6 |
| Wfdc17 | 0.70730274 | 4.17E-94 | 6 |
| Lgmn | 0.68393924 | 1.05E-11 | 6 |
| Ms4a7 | 0.67911944 | 5.19E-64 | 6 |
| Ccl4 | 0.67813758 | 5.00E-17 | 6 |
| Spi1 | 0.67520523 | 1.83E-290 | 6 |
| Mpeg1 | 0.67026454 | 1.33E-95 | 6 |
| Cebpb | 0.66528578 | 5.56E-34 | 6 |
| Hmox1 | 0.66321685 | 2.18E-08 | 6 |
| Arhgdib | 0.66236133 | 4.49E-28 | 6 |
| Itgb2 | 0.65956344 | 1.20E-185 | 6 |
| Clec4e | 0.65638449 | 9.46E-133 | 6 |
| Trem2 | 0.64565305 | 1.25E-83 | 6 |
| Fcgr2b | 0.63586868 | 1.64E-87 | 6 |
| Fcgr3 | 0.63451839 | 2.75E-220 | 6 |
| H2-D1 | 0.63154167 | 1.73E-43 | 6 |
| Plek | 0.63104102 | 5.10E-100 | 6 |
| Ftl1 | 0.63094588 | 1.37E-45 | 6 |
| Clec4d | 0.61975007 | 4.34E-137 | 6 |
| Ctsd | 0.6082818 | 3.52E-15 | 6 |
| Fth1 | 0.60104106 | 3.43E-47 | 6 |
| Lilrb4a | 0.58777686 | 5.36E-136 | 6 |
| Ptprc | 0.58248019 | 1.99E-180 | 6 |
| Csf1r | 0.57926678 | 3.68E-67 | 6 |
| Gmfg | 0.57361768 | 1.29E-228 | 6 |
| Ctsb | 0.56947148 | 2.29E-15 | 6 |
| H2-K1 | 0.56082159 | 4.30E-16 | 6 |
| Tmsb4x | 0.55987559 | 1.71E-85 | 6 |
| Cd83 | 0.55584779 | 6.12E-55 | 6 |
| Ncf2 | 0.55175718 | 1.77E-241 | 6 |
| Cd53 | 0.54967145 | 1.39E-150 | 6 |
| Cotl1 | 0.5448102 | 4.96E-27 | 6 |
| Itgam | 0.54391737 | 1.87E-176 | 6 |
| Gpx1 | 0.5416996 | 4.09E-23 | 6 |

| | | | |
|---|---|---|---|
| Lilr4b | 0.54131351 | 3.08E-115 | 6 |
| Selplg | 0.53604425 | 6.79E-234 | 6 |
| C5ar1 | 0.52590356 | 3.20E-204 | 6 |
| Clec4n | 0.5244365 | 2.93E-65 | 6 |
| Ncf1 | 0.52140625 | 1.99E-197 | 6 |
| Hmgb2 | 0.52044561 | 3.76E-13 | 6 |
| Cfp | 0.51969301 | 4.05E-115 | 6 |
| Pim1 | 0.51959353 | 5.42E-17 | 6 |
| Ms4a6d | 0.51935316 | 6.68E-119 | 6 |
| Mcl1 | 0.50965672 | 7.93E-24 | 6 |
| Adam8 | 0.50792524 | 1.87E-78 | 6 |
| Ly6c2 | 0.50661945 | 1.19E-93 | 6 |
| Ccl3 | 0.49915953 | 9.06E-29 | 6 |
| Msrb1 | 0.48872163 | 3.87E-19 | 6 |
| Sirpa | 0.48866829 | 1.19E-17 | 6 |
| Cd300c2 | 0.48760461 | 7.50E-140 | 6 |
| Cd36 | 0.48703758 | 1.12E-46 | 6 |
| Spp1 | 0.48533948 | 2.07E-09 | 6 |
| Ms4a6c | 0.4836459 | 9.09E-101 | 6 |
| Cd93 | 0.47472541 | 1.93E-93 | 6 |
| Lst1 | 0.47253335 | 5.72E-70 | 6 |
| Mrc1 | 0.47241707 | 1.24E-31 | 6 |
| Pirb | 0.47044291 | 9.00E-215 | 6 |
| Mxd1 | 0.46816921 | 2.16E-43 | 6 |
| F13a1 | 0.46121281 | 6.30E-47 | 6 |
| Dusp1 | 0.46038971 | 1.65E-12 | 6 |
| Ncf4 | 0.45985326 | 3.67E-216 | 6 |
| Bcl2a1b | 0.45982458 | 3.37E-156 | 6 |
| Lrg1 | 0.45378145 | 1.49E-143 | 6 |
| Ly6g | 0.45124535 | 1.08E-155 | 6 |
| Vsir | 0.44975716 | 1.45E-44 | 6 |
| Mafb | 0.44811273 | 1.38E-05 | 6 |
| Gda | 0.44579197 | 8.55E-48 | 6 |
| Btg1 | 0.44302046 | 4.25E-13 | 6 |
| Ctsc | 0.44258539 | 9.51E-14 | 6 |
| Taldo1 | 0.44200992 | 6.42E-16 | 6 |
| Hdc | 0.4397454 | 1.26E-111 | 6 |
| Tnf | 0.43954601 | 5.49E-66 | 6 |
| Unc93b1 | 0.4379162 | 6.54E-18 | 6 |
| Csf2ra | 0.43716002 | 8.23E-70 | 6 |
| Fyb | 0.43670302 | 6.91E-164 | 6 |
| Atp6v0c | 0.43203404 | 5.31E-22 | 6 |

| | | | |
|---|---|---|---|
| B2m | 0.43123918 | 4.15E-17 | 6 |
| Tpd52 | 0.43069859 | 3.64E-49 | 6 |
| Il1rn | 0.43060932 | 4.53E-53 | 6 |
| Pltp | 0.42858397 | 0.02003919 | 6 |
| Aif1 | 0.4253287 | 4.56E-107 | 6 |
| Grn | 0.42473934 | 1.02E-06 | 6 |
| Ptpn6 | 0.42436498 | 4.44E-224 | 6 |
| Ifi30 | 0.42052425 | 0.00227324 | 6 |
| Ptpn18 | 0.41640764 | 1.86E-125 | 6 |
| Pla2g7 | 0.41464162 | 2.21E-78 | 6 |
| G0s2 | 0.41250931 | 6.70E-16 | 6 |
| Ccrl2 | 0.41153209 | 1.04E-38 | 6 |
| Gm2a | 0.40343975 | 4.07E-08 | 6 |
| Tlr2 | 0.40289809 | 2.10E-52 | 6 |
| Efhd2 | 0.40104252 | 1.64E-11 | 6 |
| Cd177 | 0.40075389 | 2.90E-149 | 6 |
| C3ar1 | 0.39985566 | 1.80E-82 | 6 |
| Cxcr4 | 0.39880238 | 6.31E-89 | 6 |
| Nlrp3 | 0.39473639 | 9.77E-75 | 6 |
| Fabp5 | 0.39232374 | 4.92E-09 | 6 |
| Fermt3 | 0.39163226 | 1.14E-152 | 6 |
| Cdk2ap2 | 0.37346563 | 5.96E-08 | 6 |
| Ccr1 | 0.37047747 | 2.85E-223 | 6 |
| Cd24a | 0.36892618 | 2.64E-75 | 6 |
| Pld4 | 0.3683292 | 1.10E-87 | 6 |
| Osm | 0.36440135 | 5.09E-145 | 6 |
| Cytip | 0.36422853 | 3.24E-116 | 6 |
| Gpnmb | 0.36218954 | 0.00108799 | 6 |
| Ifi27l2a | 0.36193925 | 1.54E-12 | 6 |
| Mcemp1 | 0.35880184 | 2.29E-162 | 6 |
| Adgre1 | 0.35847933 | 1.26E-73 | 6 |
| Lpl | 0.35705175 | 6.15E-10 | 6 |
| Gpsm3 | 0.35310309 | 8.74E-42 | 6 |
| Arrb2 | 0.35054377 | 1.17E-75 | 6 |
| Slc7a11 | 0.34698826 | 1.20E-106 | 6 |
| Cstb | 0.34524344 | 0.00144151 | 6 |
| Ets2 | 0.34152151 | 2.66E-12 | 6 |
| Cd84 | 0.34117368 | 3.24E-188 | 6 |
| Gngt2 | 0.34108958 | 3.06E-07 | 6 |
| Lyn | 0.34074416 | 8.61E-50 | 6 |
| Atp6v0b | 0.33777636 | 6.95E-06 | 6 |
| Clec4a2 | 0.33776685 | 1.27E-215 | 6 |

| | | | |
|---|---|---|---|
| Tspo | 0.33722748 | 1.75E-13 | 6 |
| Il10 | 0.33465177 | 6.26E-09 | 6 |
| Ly86 | 0.33290878 | 3.62E-95 | 6 |
| Cxcl16 | 0.33250376 | 1.25E-09 | 6 |
| Plac8 | 0.33181529 | 1.68E-05 | 6 |
| Ubl3 | 0.33050319 | 6.99E-08 | 6 |
| Myo1f | 0.33011914 | 7.84E-225 | 6 |
| Arg1 | 0.32922296 | 4.35E-09 | 6 |
| Sat1 | 0.32877132 | 3.37E-05 | 6 |
| Slc11a1 | 0.3283651 | 6.14E-130 | 6 |
| Plin2 | 0.3282528 | 1.95E-06 | 6 |
| Ccr2 | 0.32793924 | 9.75E-47 | 6 |
| Samsn1 | 0.32557664 | 3.65E-179 | 6 |
| Card19 | 0.32341332 | 1.42E-08 | 6 |
| Rgs1 | 0.32303742 | 5.02E-44 | 6 |
| Limd2 | 0.32125164 | 7.44E-11 | 6 |
| Hck | 0.31772614 | 1.08E-174 | 6 |
| Iqgap1 | 0.31692155 | 1.46E-10 | 6 |
| Atp6v1b2 | 0.31593725 | 0.02543729 | 6 |
| H2afz | 0.31448667 | 5.25E-06 | 6 |
| Fcrls | 0.31393246 | 5.52E-45 | 6 |
| Clec12a | 0.31186586 | 7.58E-132 | 6 |
| Grina | 0.30753852 | 1.89E-06 | 6 |
| Ninj1 | 0.30619049 | 0.0036273 | 6 |
| Inpp5d | 0.30297733 | 6.40E-176 | 6 |
| Creg1 | 0.30243134 | 0.02540132 | 6 |
| Rgs2 | 0.30097042 | 3.45E-21 | 6 |
| Sdcbp | 0.30093308 | 0.00192883 | 6 |
| Plaur | 0.30004044 | 2.71E-07 | 6 |
| Hcls1 | 0.29986642 | 5.84E-256 | 6 |
| Tnfrsf1b | 0.29921973 | 4.06E-26 | 6 |
| Tnfsf9 | 0.29871043 | 3.80E-09 | 6 |
| Nrros | 0.29622944 | 1.34E-92 | 6 |
| Fcgr1 | 0.29592725 | 7.38E-123 | 6 |
| Plbd1 | 0.29370188 | 3.73E-156 | 6 |
| Pycard | 0.29325135 | 5.72E-10 | 6 |
| Slc15a3 | 0.2919037 | 1.23E-137 | 6 |
| Cd300a | 0.29165204 | 1.60E-163 | 6 |
| Snx20 | 0.29131115 | 8.06E-178 | 6 |
| Il1r2 | 0.28881466 | 1.37E-37 | 6 |
| Prkcd | 0.28850106 | 1.55E-08 | 6 |
| Fam49b | 0.28146193 | 2.23E-17 | 6 |

| | | | |
|---|---|---|---|
| Msr1 | 0.28062515 | 1.27E-74 | 6 |
| Cd300lf | 0.27962869 | 4.54E-144 | 6 |
| Nfkbia | 0.27899449 | 2.06E-07 | 6 |
| Plekho1 | 0.2782171 | 3.51E-05 | 6 |
| Capg | 0.27704146 | 2.13E-07 | 6 |
| Junb | 0.27512606 | 5.16E-11 | 6 |
| Rhog | 0.27215235 | 1.45E-10 | 6 |
| Litaf | 0.27112125 | 2.57E-06 | 6 |
| Mpp1 | 0.27066637 | 7.15E-05 | 6 |
| S100a4 | 0.27024262 | 4.41E-08 | 6 |
| Irf7 | 0.26973736 | 3.61E-10 | 6 |
| Cyth4 | 0.26969863 | 4.56E-165 | 6 |
| Arpc3 | 0.26955401 | 2.70E-10 | 6 |
| Pygl | 0.26844738 | 4.69E-14 | 6 |
| Irf5 | 0.26802284 | 4.71E-67 | 6 |
| Tmem189 | 0.26693048 | 0.00292909 | 6 |
| Gsr | 0.2642895 | 5.65E-05 | 6 |
| Ftl1-ps1 | 0.26423937 | 0.00675295 | 6 |
| Clec4a1 | 0.2636223 | 1.02E-103 | 6 |
| Csf2rb | 0.26356698 | 9.52E-192 | 6 |
| Lamp1 | 0.26330675 | 0.03761155 | 6 |
| Dock2 | 0.26305083 | 3.45E-179 | 6 |
| Hcst | 0.26286007 | 3.28E-169 | 6 |
| Rasgef1b | 0.26282016 | 7.30E-30 | 6 |
| Cd72 | 0.26278417 | 8.36E-32 | 6 |
| Ccr5 | 0.26204419 | 4.91E-92 | 6 |
| Ctsz | 0.26195623 | 0.03101334 | 6 |
| Samhd1 | 0.26170064 | 2.62E-07 | 6 |
| Nckap1l | 0.26132654 | 1.85E-157 | 6 |
| Tnfaip8 | 0.26024674 | 2.14E-08 | 6 |
| Tnfaip2 | 0.25838785 | 8.51E-05 | 6 |
| Ptpre | 0.25802867 | 5.32E-49 | 6 |
| Nfkbid | 0.25768571 | 4.75E-64 | 6 |
| Mgst1 | 0.25519376 | 4.97E-05 | 6 |
| Pilra | 0.25476678 | 2.27E-196 | 6 |
| Dusp2 | 0.25401871 | 2.36E-29 | 6 |
| Prdx5 | 0.25326957 | 0.00183343 | 6 |
| Rasgrp2 | 0.25267255 | 2.52E-77 | 6 |
| Zfp36 | 0.25219056 | 6.79E-06 | 6 |
| Syk | 0.25085039 | 5.08E-94 | 6 |
| Stab1 | 0.25003035 | 1.13E-19 | 6 |

**Table S4.** Patient information for human specimens.

| Sample No. | Age | Gender | Location of fracture |
|---|---|---|---|
| 1 | 53 | Male | Rib |
| 2 | 17 | Female | Tibia |
| 3 | 41 | Female | Rib |

**Table S5.** Antibodies used.

| Antibody | Company | Catalog # | Use |
|---|---|---|---|
| Mouse anti-Mouse CD31 | Abcam | ab24590 | IF |
| Rat anti-Mouse CD31 | BD Pharmingen | 551262 | F |
| Rat anti-Mouse CD45 | BD Pharmingen | 559864 | F |
| Rabbit anti-Mouse CD68 | Abcam | ab125212 | IF |
| Rabbit anti-Mouse F4/80 | Abcam | ab100790 | IF |
| Rabbit anti-Mouse Ki67 | Abcam | ab16667 | IF |
| Rabbit anti-Mouse NGF | Abcam | ab6199 | IF |
| Rabbit anti-Mouse/human Osteocalcin | Abcam | ab93876 | IF |
| Mouse anti-Mouse/human p75 | Santa Cruz | sc-271708 | IF |
| Rabbit anti-Mouse p75 | Abcam | ab227509 | IF |
| Goat anti-Mouse PDGFRα | R&D Systems | AF1062 | IF |
| Rat anti-Mouse PDGFRα | BD Pharmingen | 562774 | F |
| Rabbit anti-Mouse TUBB3 | Abcam | ab18207 | IF |
| Rat anti-Mouse Ter119 | BD Pharmingen | 557909 | F |
| Rabbit anti-Mouse TNFα | Abcam | ab6671 | IF |
| Donkey anti-Goat AF647 | Abcam | ab150135 | IF |
| Goat anti-Mouse AF647 | Abcam | ab150119 | IF |
| Goat anti-Rabbit AF647 | Abcam | ab150079 | IF |
| Goat anti-Rabbit IgG(H+L), DyLight 594 | Vector Laboratories | DI-1594 | IF |
| F: Flow cytometry; IF: Immunofluorescent staining. | | | |

**Table S6.** RT-PCR primers used.

| Genes (Mouse) | Forward | Reverse |
|---|---|---|
| *Bglap* | 5’-AAGCAGGAGGGCAATAAGGT-3’ | 5’-TTTGTAGGCGGTCTTCAAGC-3’ |
| *Col1a1* | 5’-TGTGTGCGATGACGTGCAAT-3’ | 5’-GGGTCCCTCGACTCCTACA-3’ |
| *Fos* | 5’-CGGGTTTCAACGCCGACTA-3’ | 5’-TTGGCACTAGAGACGGACAGA-3’ |
| *Gapdh* | 5’-GACTTCAACAGCAACTCCCAC-3’ | 5’-TCCACCACCCTGTTGCTGTA-3’ |
| *Mapk8* | 5’-AGCAGAAGCAAACGTGACAAC-3’ | 5’-GCTGCACACACTATTCCTTGAG-3’ |
| *Mapk10* | 5’-AGGTGGACAACCAGTTCTACA-3’ | 5’-GCACAGACTATTCCCTGAGCC-3’ |
| *Nfkb1* | 5’-ATGGCAGACGATGATCCCTAC-3’ | 5’-TGTTGACAGTGGTATTTCTGGTG-3’ |
| *Nfkb2* | 5’-GGCCGGAAGACCTATCCTACT-3’ | 5’-CTACAGACACAGCGCACACT-3’ |
| *Ngf* | 5’-CACTGCTCTACACCCACCCA-3’ | 5’-AGGCAGCCACAGGGGAATAG-3’ |
| *Ngfr* | 5’-CCATCTTGGCTGCTGTGGTT-3’ | 5’-GCTGTTCCATCTCTTGAAAGCAA-3’ |
| *Rela* | 5’-AGGCTTCTGGGCCTTATGTG-3’ | 5’-TGCTTCTCTCGCCAGGAATAC-3’ |
| *Relb* | 5’-CCGTACCTGGTCATCACAGAG-3’ | 5’-CAGTCTCGAAGCTCGATGGC-3’ |
| *Rhoa* | 5’-AGCTTGTGGTAAGACATGCTTG-3’ | 5’-GTGTCCCATAAAGCCAACTCTAC-3’ |
| *Rock1* | 5’-GACTGGGGACAGTTTTGAGAC-3’ | 5’-GGGCATCCAATCCATCCAGC-3’ |
| *Rock2* | 5’-TTGGTTCGTCATAAGGCATCAC-3’ | 5’-TGTTGGCAAAGGCCATAATATCT-3’ |
| *Runx2* | 5’-TGTTCTCTGATCGCCTCAGTG-3’ | 5’-CCTGGGATCTGTAATCTGACTCT-3’ |

| Genes (Human) | Forward | Reverse |
|---|---|---|
| *ALP* | 5’-ACCACCACGAGAGTGAACCA-3’ | 5’-CGTTGTCTGAGTACCAGTCCC-3’ |
| *BGLAP* | 5’-CACTCCTCGCCCTATTGGC-3’ | 5’-CCCTCCTGCTTGGACACAAAG-3’ |
| *GAPDH* | 5’-CTGGGCTACACTGAGCACC-3’ | 5’-AAGTGGTCGTTGAGGGCAATG-3’ |
| *NGFR* | 5’-CCTACGGCTACTACCAGGATG-3’ | 5’-CACACGGTGTTCTGCTTGT-3’ |
| *SP7* | 5’-CCTCTGCGGGACTCAACAAC-3’ | 5’-AGCCCATTAGTGCTTGTAAAGG-3’ |

**Table S7.** Morphogen pathway gene list.

| **Notch** | **Hippo** | **FGF** | **TGFb** | **BMP** |
|---|---|---|---|---|
| Adam17 | Yap1 | Ccn2 | Cdkn2b | Acvr1 |
| Aph1a | Wwtr1 | Cep57 | Crebbp | Amh |
| Aph1b | Ppp2ca | Crkl | E2f4 | Bmp2 |
| Aph1c | Ppp2cb | Ctnnb1 | E3f5 | Bmp4 |
| Crebbp | Ppp2r2c | Dstyk | Ep300 | Bmp5 |
| Dll1 | Ppp2r1a | Fam20c | Rbl1 | Bmp6 |
| Dll3 | Ppp2r2d | Fat4 | Smad2 | Bmp7 |
| Dll4 | Ppp2r2a | Fgf1 | Smad3 | Bmp8a |
| Dtx1 | Ppp2r2b | Fgf10 | Smad4 | Bmpr1a |
| Dtx2 | Ppp2r1b | Fgf11 | Sp1 | Bmpr1b |
| Dtx3 | Rassf6 | Fgf12 | Tfdp1 | Bmpr2 |
| Dtx3l | Wtip | Fgf13 | Tgfb1 | Gdf5 |
| Dtx4 | Ajuba | Fgf14 | Tgfb2 | Gdf6 |
| Ep300 | Limd1 | Fgf15 | Tgfb3 | Gdf7 |
| Hes1 | Trp73 | Fgf16 | Tgfbr1 | Hamp |
| Hes5 | Tead1 | Fgf17 | Tgfbr2 | Hamp2 |
| Hey1 | Tead2 | Fgf18 | Thbs1 | Id1 |
| Jag1 | Tead3 | Fgf2 | Fstl1 | Id2 |
| Jag2 | Tead4 | Fgf20 | | Id3 |
| Kat2a | | Fgf21 | | Id4 |
| Kat2b | | Fgf22 | | Inhbb |
| Lfng | | Fgf23 | | Rgma |
| Maml1 | | Fgf3 | | Rgmb |
| Maml2 | | Fgf4 | | Smad1 |
| Maml3 | | Fgf5 | | Smad4 |
| Mfng | | Fgf6 | | Smad5 |
| Ncstn | | Fgf7 | | Smad9 |
| Notch1 | | Fgf8 | | |
| Notch2 | | Fgf9 | | |
| Notch3 | | Fgfbp1 | | |
| Notch4 | | Fgfbp3 | | |
| Psen1 | | Fgfr1 | | |
| Psen2 | | Fgfr2 | | |
| Psenen | | Fgfr3 | | |
| Ptcra | | Fgfr4 | | |
| Rbpj | | Fgfrl1 | | |
| Rbpjl | | Flrt1 | | |
| Rfng | | Flrt2 | | |
| Snw1 | | Flrt3 | | |

| | | |
|---|---|---|
| | | Frs2 |
| | | Frs3 |
| | | Grb2 |
| | | Hhip |
| | | Iqgap1 |
| | | Kif16b |
| | | Kl |
| | | Klb |
| | | Lrit3 |
| | | Ndst1 |
| | | Nog |
| | | Nrxn1 |
| | | Prkd2 |
| | | Ptpn11 |
| | | Rab14 |
| | | Setx |
| | | Shcbp1 |
| | | Smoc2 |
| | | Sos1 |
| | | Sulf1 |
| | | Trim71 |

**Table S8.** Inflammatory pathway gene list.

| TNFα | IL1 | IL6 | Complement | TLR |
|---|---|---|---|---|
| Akt1 | Il1a | Il6 | C1qa | Akt1 |
| Akt2 | Il1b | Il10 | C1qb | Akt2 |
| Akt3 | Il1r1 | Il10rb | C1qc | Akt3 |
| Atf2 | Il1rap | Il10ra | C1ra | Casp8 |
| Atf4 | Myd88 | Jak1 | C1rb | Fadd |
| Atf6b | Irak1 | Jak2 | C1s1 | Il12a |
| Birc2 | Irak2 | Jak3 | C1s2 | Il12b |
| Birc3 | Irak4 | Tyk2 | C2 | Il1b |
| Casp3 | Map3k7 | Stat1 | C3 | Il6 |
| Casp7 | Traf6 | Stat2 | C4a | Myd88 |
| Casp8 | Tab1 | Stat3 | C4b | Nfkb1 |
| Cebpb | Tab2 | Stat4 | | Pik3ca |
| Chuk | Ikbkb | Stat5a | | Pik3cb |
| Creb1 | Ikbke | Stat5b | | Pik3cd |
| Creb3 | Nfkb1 | Stat6 | | Pik3r1 |
| Creb3l1 | Map2k1 | Irf9 | | Pik3r2 |
| Creb3l2 | Jnk | Crebbp | | Pik3r3 |
| Creb3l3 | Mapk9 | Ep300 | | Rac1 |
| Creb3l4 | Mapk8 | Bcl2 | | Rela |
| Creb5 | Mapk11 | Mcl1 | | Tirap |
| Dab2ip | Mapk13 | Bcl2l1 | | Tlr1 |
| Dnm1l | Mapk14 | Pim1 | | Tlr2 |
| Fadd | Mapk3 | Myc | | Tlr6 |
| Fos | Mapk4 | Ccnd1 | | Tnf |
| Ifnb1 | Jun | Ccnd2 | | Tollip |
| Ikbkb | Fos | Ccnd3 | | |
| Ikbkg | Jdp2 | Cdkn1a | | |
| Irf1 | Atf2 | Aox1 | | |
| Itch | Atf4 | Aox2 | | |
| Jun | Atf5 | Aox3 | | |
| Lta | | Aox4 | | |
| Map2k1 | | Gfap | | |
| Map2k3 | | | | |
| Map2k4 | | | | |
| Map2k6 | | | | |
| Map2k7 | | | | |
| Map3k14 | | | | |
| Map3k5 | | | | |
| Map3k7 | | | | |

| |
|---|
| Map3k8 |
| Mapk1 |
| Mapk10 |
| Mapk11 |
| Mapk12 |
| Mapk13 |
| Mapk14 |
| Mapk3 |
| Mapk8 |
| Mapk9 |
| Mlkl |
| Nfkb1 |
| Nfkbia |
| Pgam5 |
| Pik3ca |
| Pik3cb |
| Pik3cd |
| Pik3r1 |
| Pik3r2 |
| Pik3r3 |
| Rela |
| Ripk1 |
| Ripk3 |
| Rps6ka4 |
| Rps6ka5 |
| Tab1 |
| Tab2 |
| Tab3 |
| Tnf |
| Tnfrsf1a |
| Tnfrsf1b |
| Tradd |
| Traf1 |
| Traf2 |
| Traf3 |
| Traf5 |
